# DBL-1/BMP signaling pathway Smads and master transcription regulator BLMP-1/PRDM1 interact to regulate organismal traits in *Caenorhabditis elegans*

**DOI:** 10.64898/2026.09.15.751936

**Authors:** Mohammed Farhan Lakdawala, Bhoomi Madhu, Sherin Shaji, Matt Crook, Tina L. Gumienny

## Abstract

Transforming Growth Factor beta (TGF-β) signaling helps orchestrate multiple organismal traits in animals by regulating target gene expression. However, how this pathway’s transcriptional regulators, called Smads, control target gene expression to generate different traits is not well understood. Using the *C. elegans* system, we identified the master chromatin remodeler B-lymphocyte maturation protein-1 (BLMP-1, a Krüppel-like zinc finger protein homologous to human PRDM1/BLIMP1) as a partner in Smad-mediated gene expression regulation. Previous research showed BLMP-1/PRDM1 controls organismal traits that the *C. elegans* TGF-β superfamily member DBL-1 also affects. We identified genetic interactions between the DBL-1 pathway and *blmp-1* for these traits. While BLMP-1/PRDM1 and DBL-1 pathway signaling had an additive effect on body size, *blmp-1* mutants were epistatic to DBL-1 pathway mutants for male tail development, hermaphrodite gonad development, brood size, lipid levels, movement, and survival traits. We showed DBL-1 signaling transcriptionally regulates *blmp-1* expression. We also identified a physical interaction between the DBL-1 pathway Smads and BLMP-1/PRDM1 using yeast-two hybrid analyses. Using bioinformatics and qPCR analysis, we showed that Smads and BLMP-1/PRDM1 regulate expression of common downstream target genes. Our work identifies novel interactions between the DBL-1 signaling pathway and BLMP-1, two conserved major transcriptional regulators, that ultimately influence a spectrum of organismal traits. We propose a model whereby BLMP-1/PRDM1 regulates DBL-1 signaling by acting as a gatekeeper, remodeling the chromatin architecture to control the access of the Smad complex to target genes. Together, our results reveal a direct link between a TGF-β signaling pathway and a PRDM1-family master transcription regulator and offer clues to how they interact to control larval and adult traits. Our work suggests a conserved mechanism of transcriptional control of developmental traits by the Smad-PRDM axis.

**Article Summary:** Regulation of cell signaling is critical for normal development and homeostasis. Transforming growth factor beta (TGF-β) signaling controls target gene expression through Smad transcription factors. We identified novel interactions between a *C. elegans* TGF-β pathway and the chromatin remodeler B-lymphocyte maturation protein-1 (BLMP-1). Smads and BLMP-1 regulate expression of common downstream target genes that ultimately influence a spectrum of organismal traits, including body size, tissues and organs shaped by cell migration, reproduction, movement, and survival. We propose that BLMP-1 regulates TGF-β signaling by acting as a gatekeeper, remodeling the chromatin architecture to regulate Smad complex access to target genes.

## Introduction

Transforming growth factor beta (TGF-β) superfamily signaling plays dose-dependent roles in the normal development of morphological traits in animals. From body organization in sponges to body size and tissue, organ, and appendage size and patterning in complex animals, TGF-β signaling contributes to multiple traits in a time-and tissue-dependent manner (Adamska et al. 2007; Serra and Chang 2003; Wu and Hill 2009). In the core canonical pathway for TGF-β signaling, extracellular ligands bind the receptor complex, which then activate intracellular Smad transcription factors. Activated Smads can then enter the nucleus and regulate expression of an extensive number of precisely targeted genes (Fei et al. 2010; Mullen et al. 2011). How TGF-β superfamily signaling spatiotemporally coordinates expression of different genes that are important for proper development of multiple traits constitutes a fundamental knowledge gap in the TGF-β signaling field.

How do Smads control expression of specific target genes at specific times and places? It has become increasingly clear that Smads, which bind DNA weakly, rely on other chromatin-binding proteins to help them regulate expression of their target genes (Massague and Wotton 2000). Chromatin remodelers that open the chromatin, like SWI/SNF family complexes, and/or pioneer transcription factors that can bind condensed chromatin, open it, and recruit other proteins, including Smads, can help Smads bind target gene promoters in a cell-specific manner (Hill, 2016). Pioneer transcription factors like Forkhead box and zinc finger transcription factors, including Krüppel-like zinc finger proteins, have been shown to interact with chromatin remodelers and Smads (Allen and Unterman 2007; Alliston et al. 2005; Bjork et al. 2010; Feng et al. 2014; Hata et al. 2000; Hohenauer and Moore 2012; Hu et al. 2007; Sato et al. 2008; Takahata et al. 2009; Warner et al. 2007; Yao et al. 2006; Zaret and Carroll 2011). Smads and their binding partners can then interact with the Mediator complex to control gene expression by RNA polymerase II transcription machinery (Hill, 2016). However, identification of the co-factors that interact with Smad complexes to regulate TGF-β pathway target genes and an understanding of their interactions is incomplete.

The roundworm *Caenorhabditis elegans* is a well-established animal system that has been used to identify and characterize conserved TGF-β signaling pathway members and regulators. The *C. elegans* genome encodes five ligand genes (compared to >40 in mammals), and their pathways still play diverse roles (Gumienny and Savage-Dunn 2013). DBL-1 (decapentaplegic/bone morphogenetic protein-like-1) is the *C. elegans* homolog of mammalian TGF-β superfamily members BMP2/4. DBL-1/BMP is required for body size and male tail development, impacts reproductive functions like brood size and reproductive span, and plays other roles in development, homeostasis, stress responses, and aging (Table S1) (Arneaud et al. 2022; Madhu et al. 2023; Madhu et al. 2019; Mochii et al. 1999; Morita et al. 1999). DBL-1/BMP activates a conserved receptor complex, which then activates transcription regulators called Smads (including receptor-regulated R-Smads SMA-2 and SMA-3 and common co-Smad SMA-4). The activated Smad complex can then enter the nucleus to control target gene expression. Clear traits and this conserved pathway make *C. elegans* an attractive system to identify and characterize new regulators and interactors of TGF-β signaling, including what controls Smad access to their target genes (Gumienny and Savage-Dunn 2013; Savage-Dunn and Padgett 2017).

We previously identified a highly conserved transcriptional regulator, BLMP-1, that regulates DBL-1/BMP signaling in *C. elegans* (Lakdawala et al. 2019). BLMP-1, a homolog of mammalian B lymphocyte-induced maturation protein-1 (BLIMP1/PRDM1), is one of two *C. elegans* members of the PRDI-BF1 and RIZ homology domain containing (PRDM) family of master chromatin regulators. The PRDM family includes 19 human members, whose primary molecular function in metazoans is to modify chromatin structure, either directly or by recruiting other proteins (Huang et al. 2014; John and Garrett-Sinha 2009; Turner et al. 1994). BLMP-1/PRDM1 homologs bind chromatin-modifying proteins with their N-terminal PRDI-BF1-RIZ1 homologous region (PR) domain, and bind DNA with their five C-terminal Krüppel-type zinc finger domains. Smads are known to bind PRDM3 and PRDM16, which are in a different subfamily from BLIMP1/PRDM1 with two sets of zinc finger domains, and other more distantly related transcription factors with Krüppel-type zinc finger domains (Krüppel-like factors (KLFs)) (Takahata et al. 2009). In mammals, BLIMP1/PRDM1 regulates development of limbs, heart, and sensory organs. It is also important for proper differentiation of immune cells and specification of germ cell fates (Bikoff et al. 2009; Fairfax et al. 2007; Fang et al. 2018; Robertson et al. 2007). In mammalian embryos, expression of BLIMP1/PRDM1 follows the expression of TGF-β superfamily members BMPs 2 and 4 (Hopf et al. 2011). BLIMP1 expression is reduced in knock-out mouse strains for *Bmp4*, *Bmp8a*, or the BMP pathway transcription factor gene *Smad1* (Ohinata et al. 2005; Saitou et al. 2005). In *C. elegans*, BLMP-1/PRDM1, like DBL-1/BMP, also plays important roles in development, including body length determination, cuticle morphology, and gonad formation. BLMP-1/PRDM1 also acts in stress responses (oxidative stress and cues that promote entry into the dauer diapause, an alternative third larval stage animals can enter in response to reproductively unfavorable environmental conditions) and reproduction (Table S1) (Horn et al. 2014; Huang et al. 2014; Zhang et al. 2012). BLMP-1 is genetically linked to TGF-β signaling (*via* DAF-7, another TGF-β superfamily ligand) in regulating the formation of the environmental stress-resistant dauer larval stage (Hyun et al. 2016). We previously showed that *C. elegans* BLMP-1/PRDM1 is required for the long body size caused by increased DBL-1/BMP pathway signaling (Lakdawala et al. 2019). However, little is known about how BLMP-1/PRDM1 interacts molecularly with TGF-β signaling in any developmental context, especially regarding how gene expression is regulated.

**Table S1:** Common Phenotypes Between DBL-1/BMP *Signaling Pathway and blmp-1 Mutants*.

| Phenotypes | Loss of DBL-1/BMP pathway components | Loss of *blmp-1*/PRDM1 | References |
| --- | --- | --- | --- |
| body size | small | small | (Mochii et al. 1999; Morita et al. 1999; Zhang et al. 2012) |
| male tail morphology | Lep (under-retracted) | Lep or Ore (over-retracted) | (Nelson et al. 2011) |
| gonad migration | exacerbates Mig in *unc-5(-)* background | defective | (Huang et al. 2014; Merz et al. 2003) |
| brood size | reduced | reduced | (Luo et al. 2009; Zhang et al. 2012) |
| lipid stores | reduced | lipid homeostasis gene expression altered | (Clark et al. 2021; Hyun et al. 2016) |
| movement | slower | slower | (Zhang et al. 2012; Yemini et al. 2013; Simmer et al. 2003; Maniere et al. 2011) |
| permeability to drugs | increased | increased | (Schultz et al. 2014; Sandhu et al. 2021) |
| cuticle ultrastructure | annulae width reduced, basal and medial cuticle layers disrupted | annulae and alae reduced | (Zhang et al. 2012; Schultz et al. 2014) |
| lifespan | reduced in some reports | reduced | (Samuelson et al. 2007b; Portal-Celhay et al. 2012; Madhu et al. 2023; Horn et al. 2014) |

Because the *C. elegans* traits affected by loss of DBL-1/BMP pathway signaling or BLMP-1 overlap and because of the genetic interactions between BMP signaling and BLMP-1/PRDM1 homologs in mammals, we hypothesized that DBL-1/BMP and BLMP-1/PRDM1 interact genetically and molecularly in a spatiotemporal manner to control complex organismal traits. In this study, we characterized the genetic and molecular interactions of BLMP-1 with the DBL-1/BMP signaling pathway in *C. elegans*. We show that the DBL-1/BMP signaling pathway and BLMP-1/PRDM1 genetically interact to determine some traits, but appear to be additive for other traits. ChIP-seq mining reveals SMA-3 and BLMP-1 bind many of the same target genes, and qPCR validated these results for a subset of these target genes. The DBL-1/BMP pathway transcriptionally regulates *blmp-1* expression. Furthermore, we discovered that Smad proteins physically interact with BLMP-1/PRDM1. We propose that BLMP-1/PRDM1 opens chromatin containing Smad-responsive genes to allow Smad docking and transcriptional regulation of target genes in a cell-and temporally defined manner to control complex developmental traits, reproduction, and health and lifespan. Our findings provide insight into how TGF-β signaling is regulated at the level of gene transcription in two ways: 1) by TGF-β pathway-mediated regulation of *blmp-1* expression, and 2) by the chromatin regulator BLMP-1/PRDM1 binding the Smad complex to control target gene transcription. Our work furthers our understanding of how complex traits are determined by a Smad-PRDM axis.

## Materials and Methods

### *C. elegans* strains and maintenance

All *C. elegans* strains were maintained at 20^°^C on standard nematode growth media (NGM) with *E. coli* OP50 as their food source (Stiernagle 2006). Double and triple mutants were created using standard genetic crosses and confirmed by phenotype and PCR. Strains generated and used in this work are listed in Table S2.

The *blmp-1(tm548)* allele is an 810 bp deletion of part of exon 3 and intron 3. This deletion is predicted to produce a truncated protein containing the first 254 amino acids encoding the PRDI-BF1-RIZ1 homologous region (PR) domain, but not the nuclear localization sequence or DNA-binding zinc finger motifs, and 17 amino acids encoded by part of the remaining intron 3 sequence (National Bioresource Project, Japan). Polyclonal anti-BLMP-1 antibody staining gives little signal compared to the wild type in *blmp-1(tm548)* embryos (Huang et al. 2014).

*dbl-1(nk3)* is a null mutation caused by a deletion that removes 5595 bp of 5’ untranslated region and all but 33 bp of the *dbl-1* open reading frame encoding this TGF-β superfamily member (Schultz et al. 2014).

*sma-3(e491)* is a missense mutation (G350R in the dwarfin or Mad homology region 2 (MH2)). This amino acid variation severely reduces the function of this R-Smad gene product (Savage et al. 1996).

*sma-3(wk30)* contains a premature stop codon (R145X) and is a putative null based on its strong phenotype (Savage-Dunn et al. 2000).

**Table S2.**
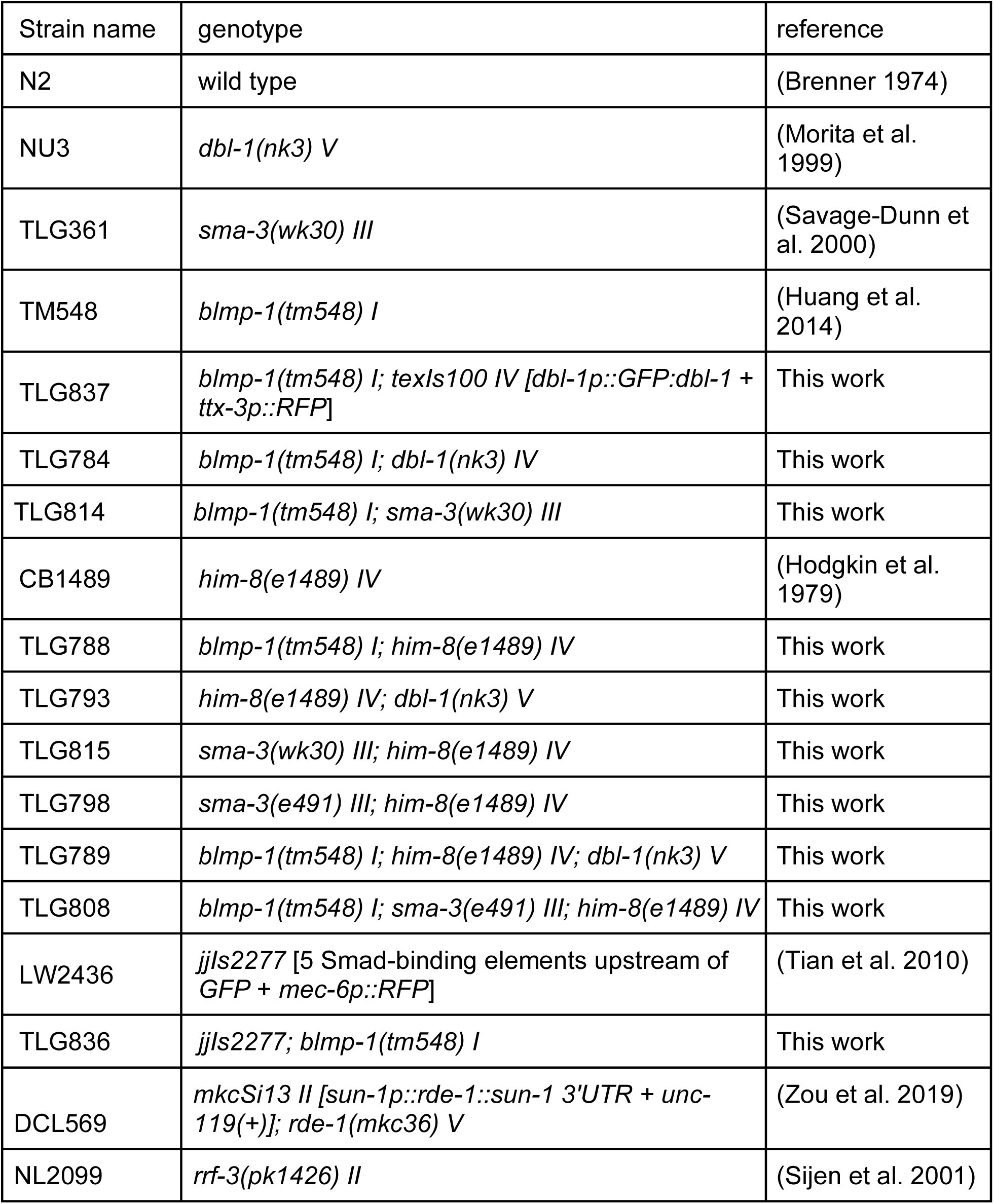
Strains used.

### Body size measurement

Body size measurements were performed as previously described (Lakdawala et al. 2019). Animals were bleach-synchronized and grown at 20^°^C on NGM plates with *E. coli* OP50. After 48 hours, L4 animals were picked on a fresh plate and allowed to grow for an additional 24 hours. Adults were mounted in 1 mM levamisole on a 2% agarose pad on a glass slide. Images were captured using a Nikon SMZ1500 dissecting microscope and body size was measured using the length measurement image tool on the iVision-Mac software. Data was statistically analyzed by one-way ANOVA using Tukey’s multiple comparisons test.

### Male tail and DTC migration

Animals were bleach-synchronized and grown to the appropriate stage. Male tail defects were scored and compared in one-day old adults as previously described (Nelson et al. 2011). Distal tip cell (DTC) migration defects were scored and compared in L4 animals as previously described (Huang et al. 2014). Animals were mounted in 1 mM levamisole on a 2% agarose pad on a glass slide and scored using Nomarski differential interference contrast (DIC) microscopy with a 40X objective on a Zeiss LSM 900 confocal system.

### Quick Oil Red O total lipid staining

Total lipid staining of individual strains was carried out using the Quick Oil Red O (qORO) method (Wählby et al. 2014). A 0.5% qORO (Alfa Aesar, MA, USA) stock solution was made in isopropanol. After an overnight incubation on a rocker at 25°C, the 0.5% qORO solution was passed through a 0.45 µm filter before being stored at room temperature. Before use, this stock solution was diluted to 0.3% in distilled water and incubated overnight before filtering a second time with a 0.45 µm filter. Mixed stage animals were washed off NGM-OP50 plates using M9 buffer, then centrifuged at 14,000 rpm for one minute, after which the supernatant was removed and replaced with 0.5 ml of isopropanol to fix the animals. Samples were immediately vortexed and centrifuged a second time. The supernatant was replaced with 0.5 ml of 0.3% qORO solution and samples were incubated overnight on a rocker at 25°C. After a brief centrifugation, the stain was removed and animals were resuspended in 250 µl of 0.1% Triton X-100 (ThermoFisher, MA, USA) in S buffer, and samples were stored at 4°C until use (Stiernagle 2006). L4 animals or young adults without eggs were picked onto 5% agar slides and imaged in whitefield at 200x total magnification with a Nikon Eclipse Ni microscope.

Each image was then processed by removing background artifacts with the “Magic Select’’ function in Microsoft Paint 3D. Images were converted using BioPython V2.7 to negative grayscale to represent the amount of light absorbed by the stain (Wang et al. 2025). The mean pixel intensity of each animal was determined in BioPython V2.7, with a higher number representing a higher level of lipid staining.

### Brood size

Brood size was performed as described previously with minor modifications (Madhu et al. 2019). NGM plates were seeded with 20 µl of bacteria a day before use and bacteria allowed to dry overnight. L4 animals were picked on individual plates. Animals were transferred to a fresh plate every 24 hours until they stopped laying eggs. The number of eggs and hatched embryos were manually counted under a dissecting microscope every 24 hours. Mean brood size was quantified and compared. Because DBL-1/BMP pathway mutant strains have a high matricide rate, all brood size experiments were started with a sample size of at least 15 animals. Animals that died before the end of the reproduction period were censored from the analyses. Data from at least seven animals per genotype was used for statistical analysis. Power analysis was done using f value of 0.8, α = 0.05, and (Power) 1-β = 0.08. Data was statistically analyzed by one-way ANOVA using Tukey’s multiple comparison test.

### RNAi experiments

RNA interference (RNAi) was performed by the feeding method as previously described (Kamath et al. 2000). Single colonies of RNAi bacteria were grown overnight in LB broth containing 50 µg/ml carbenicillin for 10–12 hours and induced to express dsRNA using 1 mM IPTG for 4–6 hours. Bacteria were grown for no more than 16 hours in total. The induced bacterial culture was seeded on NGM plates containing 50 µg/µl carbenicillin and 1 mM IPTG and dried overnight before use.

For the brood size experiments in germline-suppressed *blmp-1* animals, a mixed population of DCL569 *[sun-1p::rde-1::sun-1 3’UTR + unc-119(+)]*, a strain that allows germline-specific RNAi, was bleached and eggs were seeded on the RNAi bacteria plate. Plates were incubated at 20^°^C and animals were used for brood size experiments at the desired stage. RNAi bacteria of C06C3.5, a predicted pseudogene, was used as the negative control. For suppressing *blmp-1* only at adult stage, synchronized animals of RNAi-sensitive strain NL2099 *rrf-3(pk1426)* were grown on C06C3.5 RNAi bacteria until the L4 stage and were then transferred to *blmp-1* RNAi bacteria plates. NL2099 animals grown on *blmp-1* RNAi bacteria throughout development were used as a positive control.

### Video analyses of movement

Mid-L4 animals were placed individually on OP50 seeded NGM plates and incubated at 20°C for 24h. Each animal was separately placed on an unseeded plate and recorded at 10x magnification and 15 fps using an Olympus SZX10 dissecting microscope and DP74 camera. Each recording was approximately 1000 frames long and stored as an uncompressed .avi file. Ten to twenty animals per strain were assayed in each of five separate rounds.

Each video was analysed in ImageJ (Schneider et al. 2012) using the wrMTrck plugin (Nussbaum-Krammer et al. 2015). Briefly, each video was converted to greyscale followed by background subtraction using the rolling ball method, set to a one pixel rolling ball radius with “light background”, “sliding paraboloid” and “disable smoothing” selected. The threshold was set using “MaxEntropy” and the auto function, followed by manual adjustment for each video to minimize background noise while maintaining the integrity of the animal being imaged. Scale was set at 106 pixels per mm for all videos. We changed the following wrMTrck plugin settings from the default: minSize of 35 to 70, depending on strain, maxSize of 200, minTrackLength of 20 and FPS of 15. We averaged each strain for each round, calculated the overall average per strain for the five rounds and then normalized each strain to wild type.

### Survival analysis

Mid-L4 animals were placed individually on separate NGM-OP50 plates at 20°C and examined for movement at 24, 48 and 72h. Non-moving animals were poked with a platinum wire worm pick and scored as alive if they showed any sign of movement and dead if they failed to respond and no pharyngeal pumping was detected at the highest magnification on the dissecting microscope used. Percent survival for each day was calculated as (NUMBER OF ANIMALS ALIVE ON DAY x – NUMBER OF ANIMALS ALIVE AT DAY 0) x 100.

### DBL-1/BMP Smad reporter imaging

Double mutant animals were generated that contain *blmp-1(tm548)* and an integrated reporter containing green fluorescent protein (GFP) after five Smad binding elements (allele name *jjIs2277*, “RAD-SMAD”). Experiments with the RAD-SMAD reporter strains were performed as previously described (Savage-Dunn et al. 2019; Tian et al. 2010). Animals were bleach-synchronized and allowed to grow for 24 hours until they reached L2 stage. L2 animals were mounted in 1 mM levamisole on 2% agarose pad on a glass slide. Hypodermal nuclei GFP fluorescence of each animal was captured using 40X objective on a Zeiss LSM 900 confocal microscope. The conditions were optimized with respect to the control and were kept consistent within each trial. At least 15 animals per condition were imaged in three independent trials. Mean fluorescence intensities were measured using the ZEN 3.1 software. Data was statistically analyzed by unpaired *t*-test.

### cDNA synthesis and qRT-PCR

Animals were bleach-synchronized and grown to the desired stage. Total RNA was isolated using TRIzol™ Reagent (ThermoFisher) by freeze-crack method as described previously (Portman 2006). RNA was normalized within each trial before using it for cDNA synthesis. cDNA was synthesized using the SuperScript III reverse transcriptase kit (ThermoFisher) as per manufacturer’s protocol. cDNA was normalized within each trial before using it for real-time PCR. A real-time PCR reaction of 20µl total volume was set up using the PowerUP™ SYBR™ GREEN master mix (Applied Biosystems™) and PCR was performed using QuantStudio 3 system as per manufacturer’s instructions (Applied Biosystem). C_t_ values for each target gene were identified using the QuantStudio Design and Analysis Software and normalized to the housekeeping gene *act-1*. Fold change was determined by using the 2^(-ΔΔCt)^ method. The experiments for Figure 9 were performed in four biological replicates with three technical replicates each. The experiments for Figure 12 were performed with three technical replicates each. Data was statistically analyzed using unpaired *t*-test or one-way analysis of variance (ANOVA) using Tukey’s multiple comparisons test.

### Yeast two-hybrid

cDNA of *blmp-1* and *daf-16* were cloned as bait proteins in a pAS2 cloning vector as GAL4 DNA-binding domain (BD)-BLMP-1 and GAL4 DNA-BD-DAF-16, respectively. *sma-2*, *sma-3*, and *sma-4* cDNAs were cloned into a pACT2 vector as GAL4 activation domain (AD)-SMA-2, GAL4 AD-SMA-3, and GAL4 AD-SMA-4, respectively. DNA binding domain and activation domain plasmids were co-transformed in yeast strain AH109 using lithium acetate method (Yeastmaker™ Yeast Transformation System 2 User Manual, Clontech PT1172-1, Cat No. 630439)). Co-transformants were selected on media lacking leucine and tryptophan. To test for interaction, colonies were further screened on media lacking leucine, tryptophan, histidine, and adenine.

### Bioinformatics analyses

To visualize SMA-3 and BLMP-1 peaks, read-depth normalized signal bigwig files ENCFF290VZU (Valerie Reinke lab) for SMA-3 tagged with eGFP and ENCFF351SBS (Kevin White lab) for BLMP-1 tagged with eGFP from encodeproject.org were visualized using the Integrative genomics viewer (https://igv.org; (Robinson et al. 2011)). To identify intersections between SMA-3 and BLMP-1 peaks, the usegalaxy.org program was used with bedtools intersect intervals with default setting (The Galaxy Community, 2024; Quinlan and Hall, 2010). Optimal irreproducible discovery rate (IDR) thresholded peaks for SMA-3 (ENCFF964QGY, from the Kevin White laboratory) and BLMP-1 (ENCFF494PRR, from the Valerie Reinke laboratory) were used as input for ChIPseeker. Disease orthologs, human orthologs, and gene ontology associations in Table S3 were extracted using [SimpleMine](https://www.alliancegenome.org/agr_simplemine.cgi) through Alliance of Genome Resources.

## Results

### Loss of *blmp-1* suppresses long body size and acts independent of DBL-1/BMP signaling

The DBL-1/BMP signaling pathway is a major pathway responsible for body size regulation in *C. elegans*. The effect of DBL-1/BMP signaling is dose dependent, as an increase in signaling activity results in longer animals and a decrease in signaling makes the animals smaller in size (Gumienny and Savage-Dunn 2013; Savage-Dunn and Padgett 2017)). *blmp-1* mutants are also significantly smaller than wild-type animals (Zhang et al. 2012). Previous work showed that a *blmp-1* loss-of-function mutation (*s71*, L71*) suppresses the long body size of animals overexpressing DBL-1/BMP ligand (Lakdawala et al. 2019). To confirm if a partial deletion of *blmp-1, tm548*, can also suppress the long body size phenotype, we created a strain containing *blmp-1(tm548)* that also overexpresses DBL-1/BMP (*dbl-1(++)*) (Huang et al., 2014). *blmp-1(tm548)* significantly suppressed the long body size of DBL-1/BMP-overexpressing animals; however, the suppression was only partial when compared to *blmp-1* mutants (Figure 1). These findings suggest that functional BLMP-1/PRDM1 is necessary but not sufficient for the increased body size of animals overexpressing DBL-1/BMP. To further test if DBL-1/BMP signaling and BLMP-1/PRDM1 work together to regulate body size, we generated double mutant strains with *dbl-1(nk3)* and *blmp-1(tm548)*. We compared their body size average to that of the single mutants. Double mutants were significantly smaller than either *dbl-1* or *blmp-1* single mutants. Animals lacking both *sma-3* and *blmp-1* also displayed a smaller body size on average than the single mutant animals. These results indicate that the DBL-1/BMP pathway and BLMP-1/PRDM1 affect body size at least partly independently.

**Figure 1.**
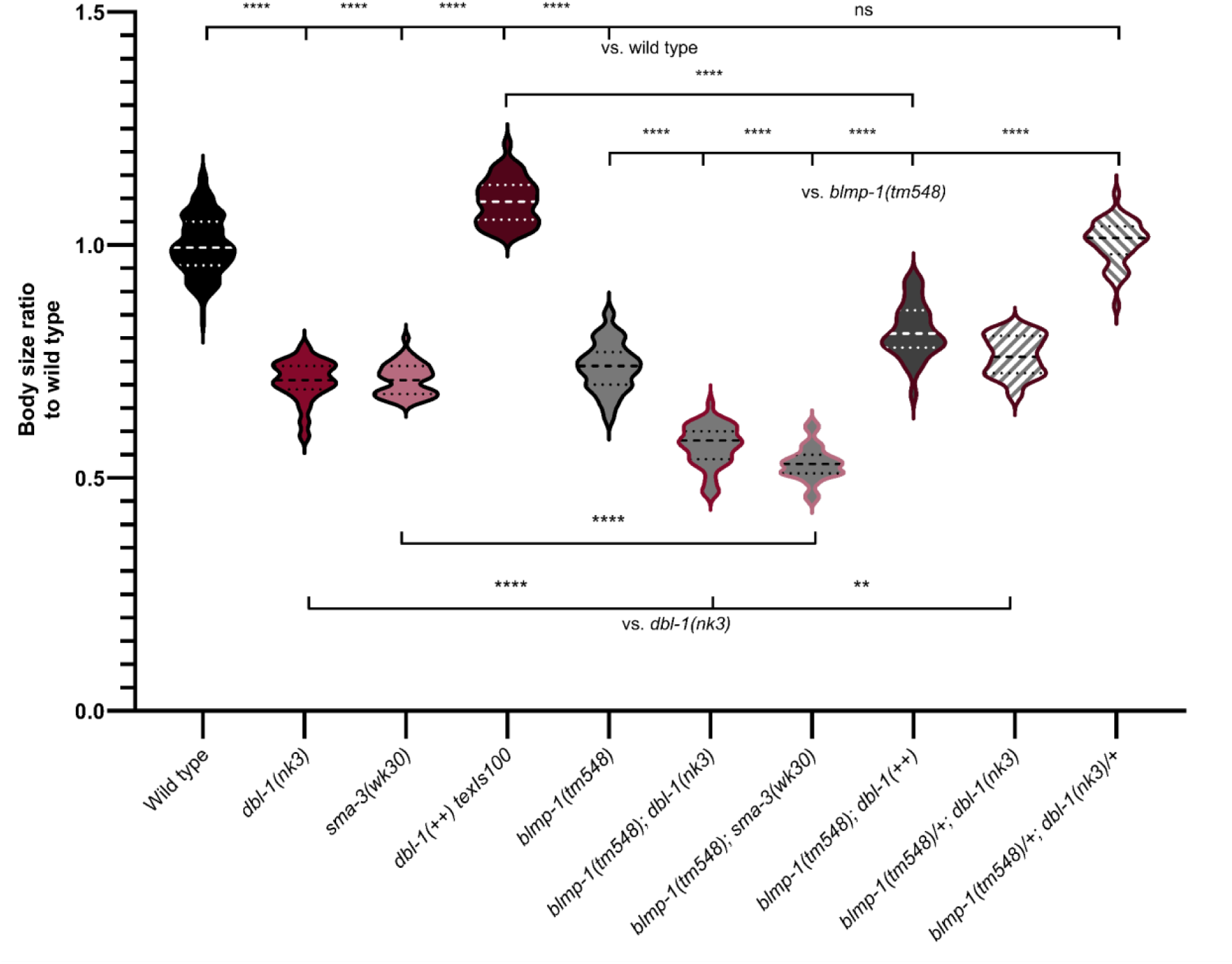
*blmp-1(–)* suppresses body size of adult animals in different DBL-1 pathway backgrounds in a recessive manner. Body size of adult animals was measured 24 hours post L4 stage. Body size of the double mutants was measured and compared to that of wild-type (WT) control or respective single mutants. n = at least 40 per condition. Error bars represent standard deviation. **** represents *p* < 0.0001, compared by one-way ANOVA using Tukey’s multiple comparisons test.

Because the partial deletion of *blmp-1* results in a visibly smaller body than the null, we asked if *blmp-1* affected body size in a dominant or a recessive manner. Animals lacking *dbl-1* but heterozygous for *blmp-1*(*tm548)* are no smaller than *dbl-1* homozygous animals (Figure 1). In the *dbl-1* heterozygous background, heterozygosity of *blmp-1*(*tm548)* had no effect on body size. The double heterozygotes were not significantly different from the wild type in body size. Together, these results indicate that *blmp-1* acts recessively downstream of DBL-1/BMP.

### *blmp-1* is epistatic to *dbl-1* for male tail morphogenesis

Defects in DBL-1/BMP signaling and BLMP-1/PRDM1 affect male tail development. In the last larval stage, the pointed tail tip retracts, generating a blunt-ended tail. Delayed or inhibited retraction results in an under-retracted, pointy adult tail (the <u>lep</u>toderan or Lep phenotype) (Nelson et al. 2011). Precocious retraction results in an <u>o</u>ver-<u>re</u>tracted, abnormally blunt tail (the Ore phenotype) (Del Rio-Albrechtsen et al. 2006). DBL-1/BMP pathway mutants have multiple male tail development defects including Lep (Nelson et al. 2011; Savage et al. 1996). In contrast, *blmp-1* animals can have either a Lep or the opposite ‘Ore’ (<u>o</u>ver-<u>re</u>tracted/abnormally blunt) tail depending on the level of *blmp-1* suppression (Emmons 2005; Nelson et al. 2011). We generated triple mutant strains with a *him-8* mutation to enrich for males. Loss of *him-8* function results in a <u>h</u>igh incidence of <u>m</u>ales with normal adult tail morphology (Figure 2, Hodgkin et al., 1979). *blmp-1(tm548)* males all had the over-retracted defect. While some males lacking *dbl-1* or *sma-3* function had normally shaped tails, the majority (averaging 56–65%) displayed the under-retracted tail defect, consistent with previously published work (Nelson et al. 2011; Savage et al. 1996). All populations with the *blmp-1(tm548); him-8(–); dbl-1(–)* or *blmp-1(tm548); him-8(–); sma-3(–)* genotype were 100% Ore, identical to *blmp-1; him-8* (Figure 2). These results indicate that *blmp-1* is epistatic to *dbl-1* and *sma-3*, with the *blmp-1(lf)* precocious retraction defect preventing the late retraction defect in DBL-1/BMP pathway mutant male tails from occurring. This result supporting a model in which BLMP-1/PRDM1 works earlier than the DBL-1/BMP pathway to regulate male tail development.

**Figure 2.**
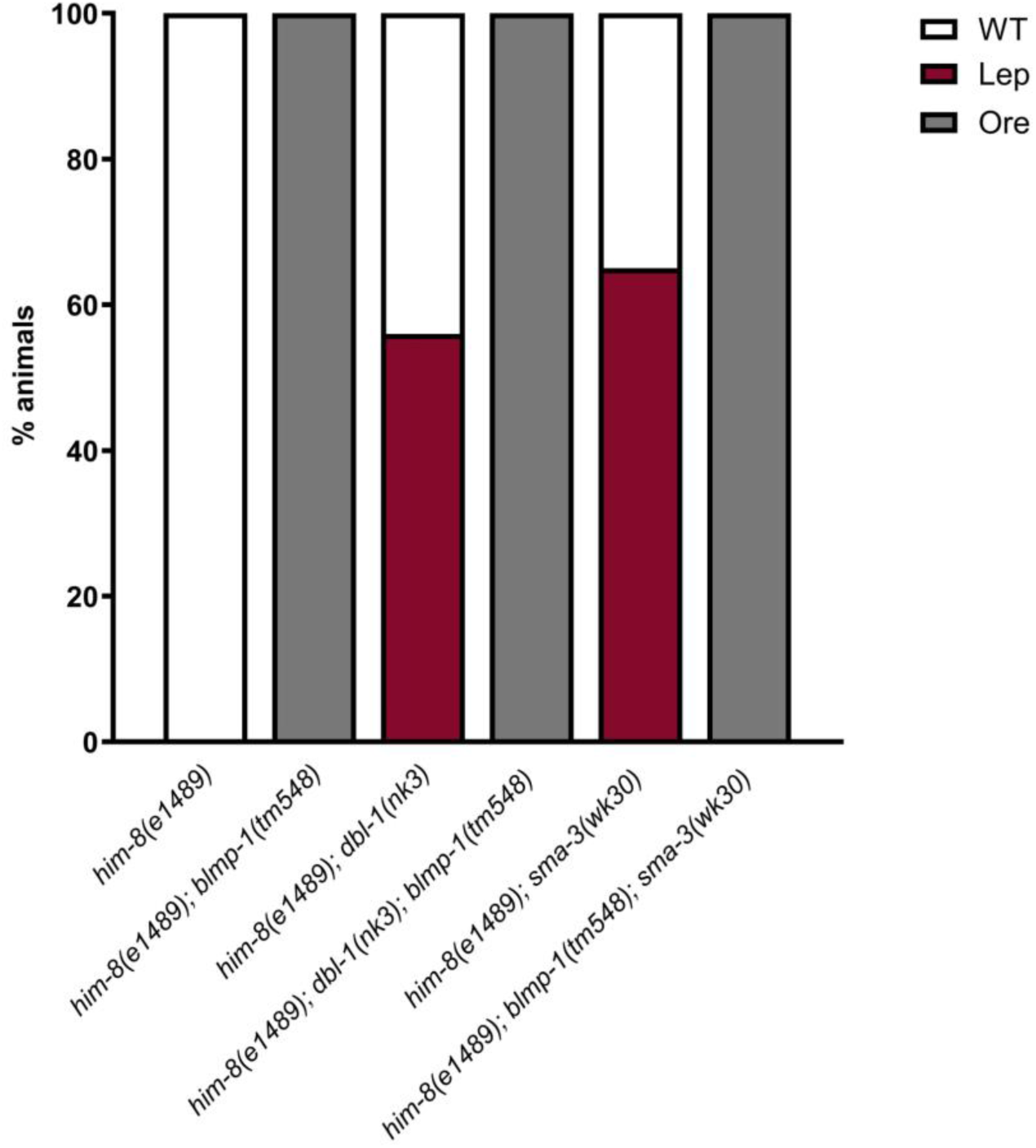
The *blmp-1(–)* over-retraction (Ore) male tail defect masks the DBL-1/BMP pathway mutant pointed/leptoderan (Lep) male tail defect. Defects in the tail morphology of *him-8*, *him-8; blmp-1(tm548)*, *him-8; dbl-1(–)*, *him-8; sma-3(–)*, *him-8; blmp-1(tm548); dbl-1(–)*, and *him-8; blmp-1(tm548); sma-3(–)* were recorded and compared 24 hours post L4 stage. The fraction of these animals exhibiting Lep, Ore, and WT phenotypes was calculated. n = at least 30 males per condition. Male tail defects of *him-8; blmp-1(tm548); dbl-1(–)* or *him-8; blmp-1(tm548); sma-3(–)* were similar to double mutant *him-8; blmp-1(tm548)*.

### DBL-1/BMP pathway mutants do not alter the *blmp-1* DTC migration defect penetrance

Normal gonad migration is a larval process that is affected by loss of DBL-1/BMP pathway and BLMP-1/PRDM1 function. UNC-5, a netrin receptor, is required for proper gonadal morphogenesis, where the two migrating distal tip cells follow netrin and other cues to each make two turns that result in two U-shaped gonads (Leung-Hagesteijn et al. 1992). Loss of *dbl-1* by itself does not have any distal tip cell (DTC) migration defects, but loss of *dbl-1* in an *unc-5* mutant background increases the penetrance of gonad migration defects caused by loss of *unc-5* (Merz et al. 2003). Reduced or overexpressed *blmp-1* levels affect *unc-5* expression levels, leading to gonad migration defects (Huang et al. 2014). We asked if loss of *dbl-1* increases the incompletely penetrant gonad migration defects of *blmp-1* mutant animals. To test the hypothesis, DTC defects were scored by scoring L4 gonad dismorphology penetrance in *blmp-1(tm548); dbl-1(–)* and *blmp-1(tm548); sma-3(–)* double mutant populations and comparing them to single mutants and the wild type. No migration defects were seen in WT, *dbl-1(–)*, or *sma-3(–)* animals (Table 1). Most, but not all, gonad arms in *blmp-1(tm548)* animals displayed defective migration. *blmp-1(tm548); dbl-1(–)* did not show significantly increased gonad migration defects when compared to *blmp-1(tm548)* animals alone (*p*-value = 0.26). Similarly, *blmp-1(tm548); sma-3(–)* animals did not exhibit increased gonad migration defects in comparison to *blmp-1* mutants (*p*-value = 0.14). These results indicate that DBL-1/BMP pathway mutants do not enhance or suppress the penetrance of the *blmp-1* DTC migration defect.

**Table 1.** The penetrance of the hermaphrodite gonad migration defect in *blmp-1(–)* is not significantly altered by loss of DBL-1/BMP signaling. Distal tip cell (DTC) migration defects were compared between wild-type (N2), *blmp-1(tm548)*, *dbl-1(nk3)*, *sma-3(wk30)*, *blmp-1(tm548); dbl-1(nk3)*, and *blmp-1(tm548); sma-3(wk30)* animals at the L4 stage. The percentages of animals demonstrating anterior and posterior DTC migration defects were calculated and statistically analyzed by Fisher’s exact test. n = at least 48 animals per group. Gonad migration defects of double mutant *blmp-1(tm548); dbl-1(nk3)* or *blmp-1(tm548); sma-3(wk30)* are not significantly different from gonad migration defects observed in *blmp-1(tm548)* single mutants.

| genotype | distal tip cell migration defect |  |  |  |
| --- | --- | --- | --- | --- |
|  | anterior | n | posterior | n |
| wild type | 0% | 100 | 0% | 100 |
| *dbl-1(nk3)* | 0% | 100 | 0% | 100 |
| *sma-3(wk30)* | 0% | 100 | 0% | 100 |
| *blmp-1(tm548)* | 88% | 135 | 77% | 132 |
| *blmp-1(tm548)*; *dbl-1(nk3)* | 70% | 53 | 58% | 48 |
| *blmp-1(tm548)*; *sma-3(wk30)* | 95% | 95 | 84% | 85 |

### BLMP-1/PRDM1 and DBL-1/BMP signaling work together to regulate brood size

DBL-1/BMP pathway mutants and *blmp-1(RNAi)* animals have a reduced brood size (Luo et al. 2010; Zhang et al. 2012). To test if there is genetic interaction between *blmp-1* and DBL-1/BMP signaling to maintain brood size, the total brood size of double mutant *blmp-1(tm548); dbl-1(nk3)* was measured and compared to single mutants and the wild-type control. Animals lacking either *dbl-1* or *sma-3* had a similarly reduced brood size (Figure 3). The average brood size of *blmp-1(tm548)* animals was fewer than the DBL-1/BMP pathway mutant brood size. The total brood size of *blmp-1(tm548)*; *dbl-1(nk3)* animals was not significantly different from the *blmp-1(tm548)* brood size. However, the *blmp-1(tm548); sma-3(wk30)* brood size average was significantly lower than either single mutant brood size. These results suggest *blmp-1* is epistatic to DBL-1/BMP signaling to regulate animal brood size.

**Figure 3.**
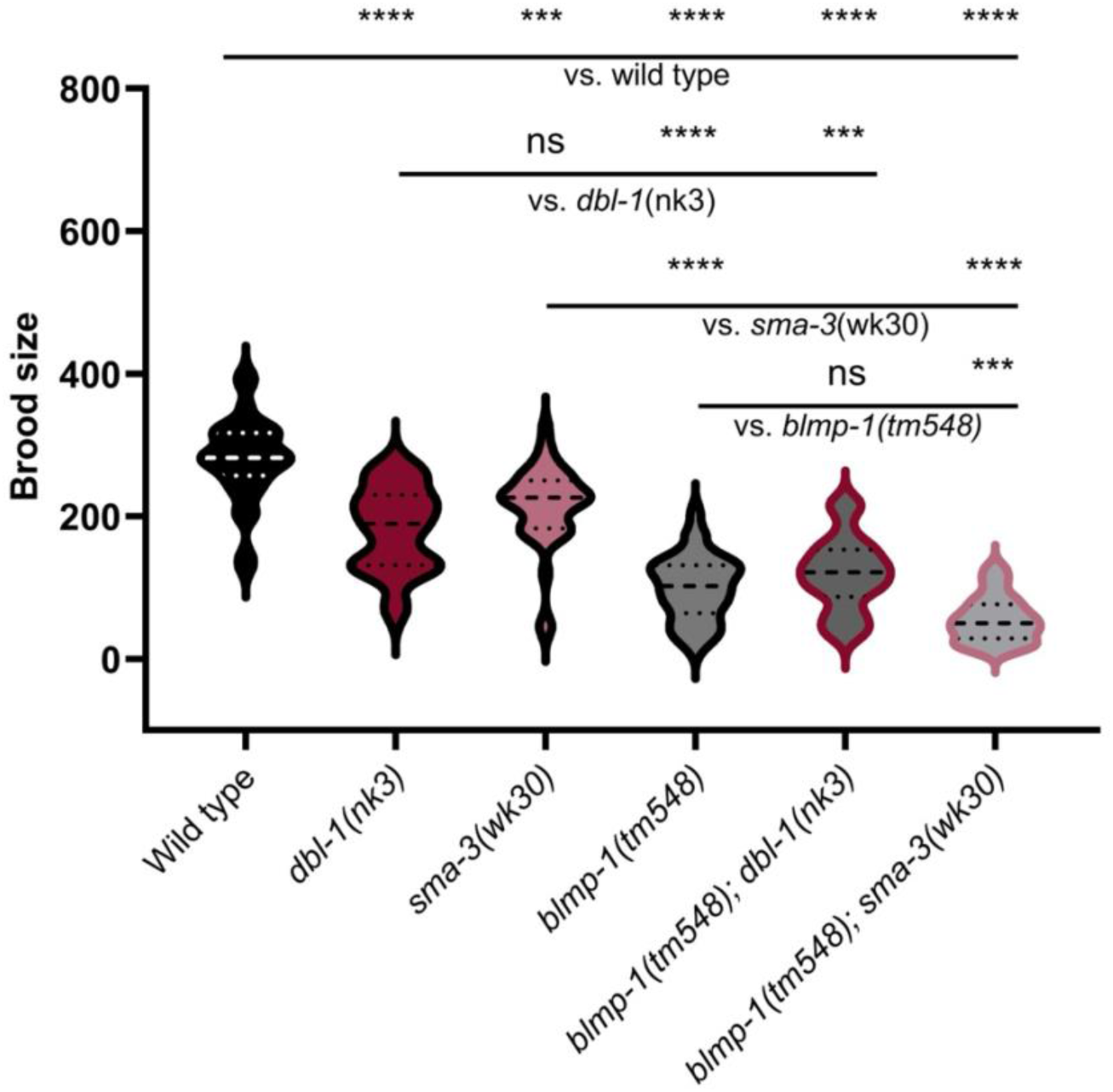
BLMP-1/PRDM1 and DBL-1/BMP signaling work together to regulate brood size. Brood size of animals picked as L4s was measured by counting the number of laid embryos and hatchlings every 24 hours until the cessation of egg laying. The average brood size of double mutants was compared to that of wild-type or respective single mutants. n = at least 7 per condition. Brood sizes less than 10 were excluded. Error bars represent standard deviation. * *p* < 0.05, ** *p* < 0.005, **** *p* < 0.0001, compared by one-way ANOVA using Tukey’s multiple comparisons test.

### BLMP-1/PRDM1 is required during development for normal brood size

Brood size of self-fertilized hermaphrodites depends on multiple factors, including somatic tissue development and germline factors. The major turns in the hermaphrodite gonad occur in L3 and early L4 stages, while the germline makes sperm in L4 and oocytes in adulthood (Lints and Hall, 2009; Lints and Hall, 2009). To identify the temporal requirement of *blmp-1* to regulate brood size, *blmp-1* was suppressed in RNAi-sensitized *rrf-3(–)* animals either throughout larval development through adulthood or from the last larval stage through adulthood. *blmp-1* suppression throughout larval development significantly reduced the animals’ brood size (Figure 4). However, *blmp-1* suppression during adulthood but not during all larval stages had no effect on the total progeny number. These results suggest wild-type brood size requires BLMP-1/PRDM1 function in gonad morphogenesis during larval development.

**Figure 4.**
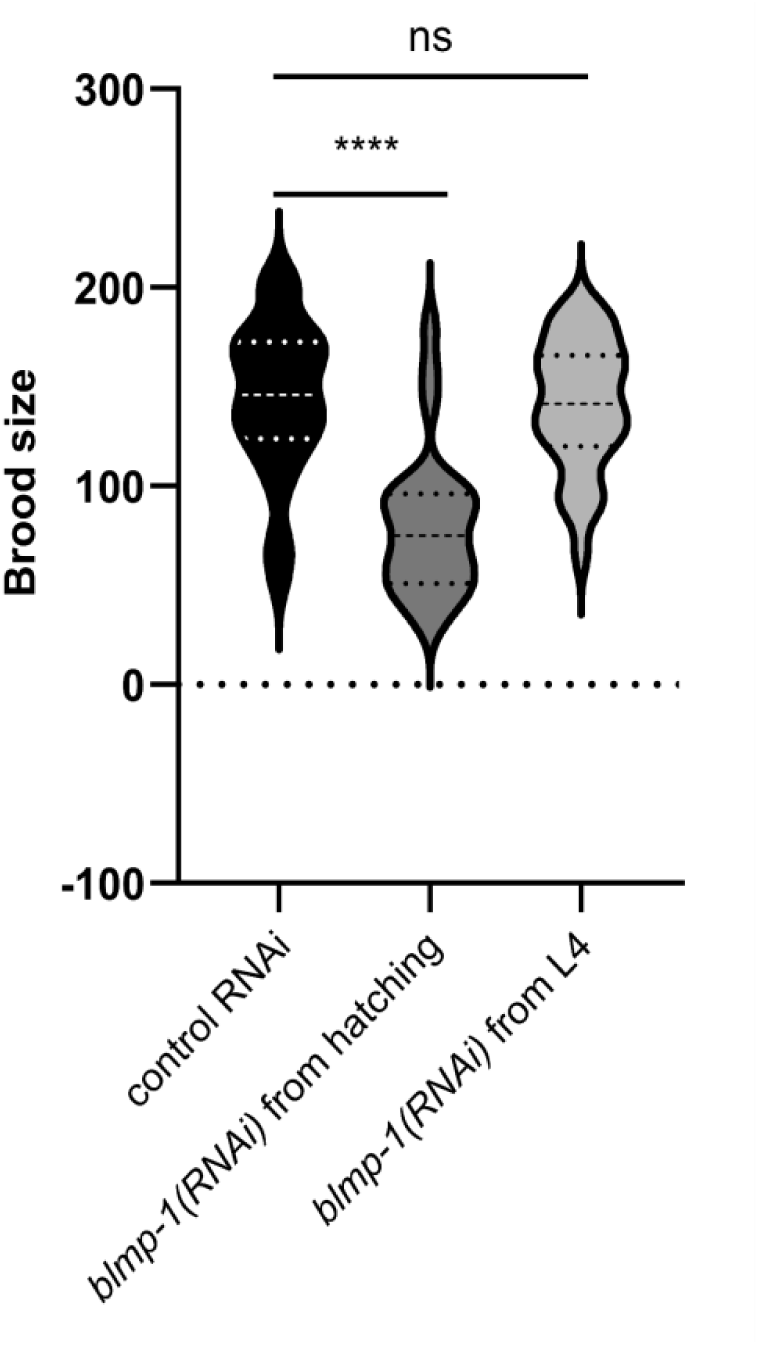
BLMP-1/PRDM1 is required during development to regulate brood size. Brood size of animals starting as embryos or picked as L4s was measured by counting the number of laid embryos and hatchlings every 24 hours until the cessation of egg laying. n = 27–30 animals per condition. Error bars represent standard deviation. *p* < 0.0001, by one-way ANOVA using Tukey’s multiple comparisons test.

### *sma-3* and *blmp-1* loss-of-function animals have lower lipid levels compared with wild-type

Loss-of-function mutations in *dbl-1* and *sma-3* have been shown to reduce lipid storage in the intestine of *C. elegans* (Clark et al. 2021). BLMP-1/PRDM1 inhibits expression of *sams-1*, an S-adenosyl methionine synthase gene that is important for lipid homeostasis (Hyun et al. 2016; Li et al. 2011). We asked if loss of BLMP-1/PRDM1 function affects lipid levels and if BLMP-1/PRDM1 interacted with the DBL-1/BMP signaling pathway for this trait. We found that *sma-3(wk30)* animals had significantly lower whole animal lipid levels compared with the wild type (93%, p<0.0001), but *dbl-1(nk3)* animals were not significantly different in our conditions (Figure 5). Notably, *blmp-1(tm548)* mutants had significantly lower lipid levels than the wild type (96%, p=0.0069).

**Figure 5.**
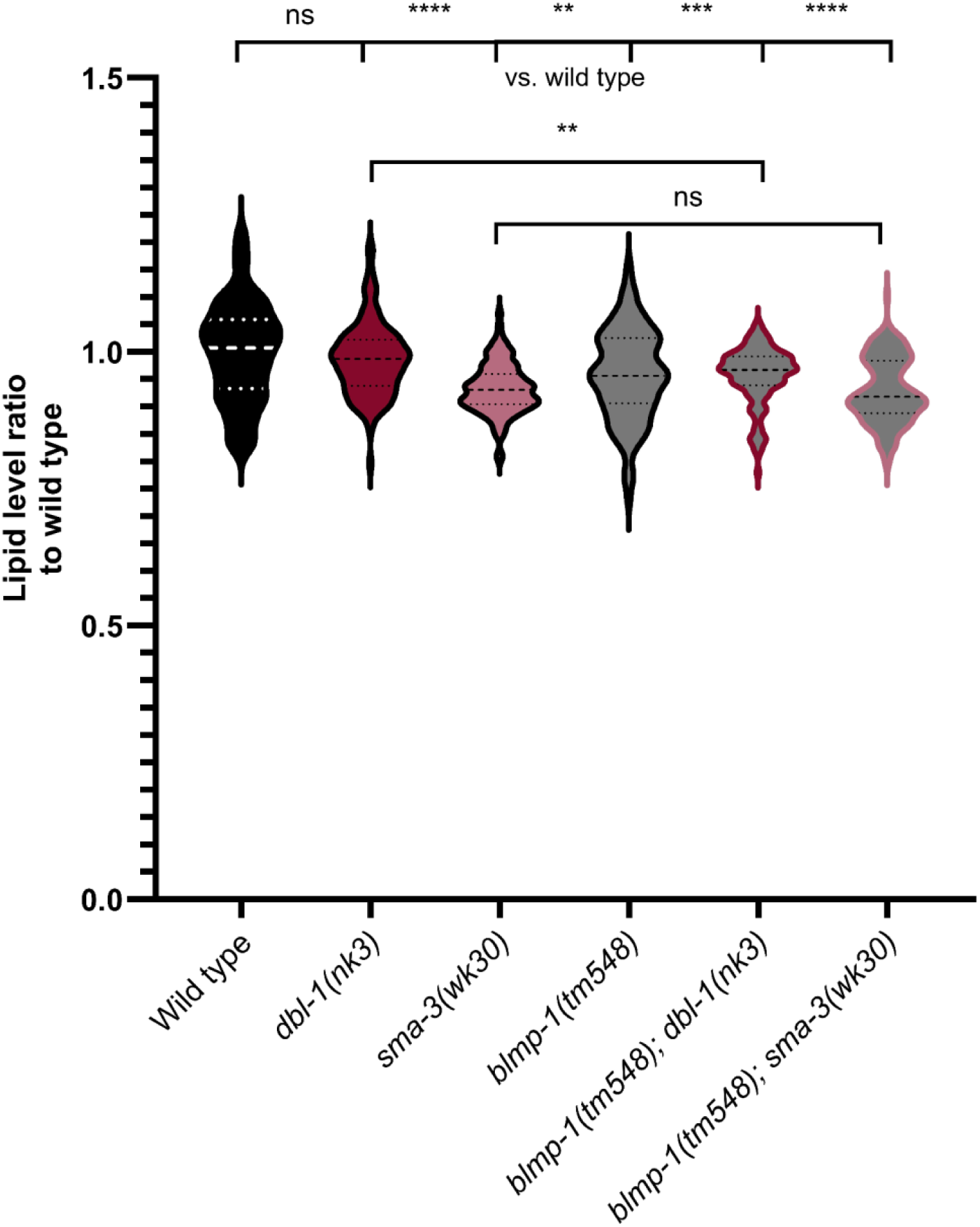
Loss of *sma-3* and *blmp-1* function reduces lipid levels compared with wild-type, but *dbl-1* loss of function has no effect. Whole animal lipid levels in L4 to young adult animals were quantified using quick Oil Red O staining for between 114 and 130 animals per genotype. In the violin plot, the central line represents the median, the lighter dotted lines show the 25^th^ and 75^th^ percentiles, respectively, and the outside line spans the full data range. The shape and width of each violin display the density of the data at different values, providing a detailed view of how data are spread within each group. ****, ***, **, * and ns represent p<0.0001, p<0.001, p<0.01, p<0.05 and non-significant, respectively, using Dunnett’s multiple comparisons tests in Welch’s ANOVAs.

To determine if a genetic interaction exists between the DBL-1/BMP signaling pathway and BLMP-1/PRDM1, we then assessed lipid levels in double mutant strains. *blmp-1(tm548); dbl-1(nk3)* mutants had significantly lower lipid levels than the *dbl-1(nk3)* mutant. The *blmp-1(tm548); sma-3(wk30)* double mutants were not significantly different from the *sma-3(wk30)* single mutant (Figure 5). These results suggest that SMA-3/R-Smad and BLMP-1/PRDM1 act in the same pathway to maintain normal lipid levels.

### *dbl-1*, *sma-3* or *blmp-1* are independently required for normal adult movement

*dbl-1* pathway and *blmp-1* mutants move slower than the wild type (Maniere et al. 2011; Simmer et al. 2003; Yemini et al. 2013; Zhang et al. 2012). We asked if there was a genetic interaction between the *dbl-1* pathway and *blmp-1* for movement. To do so, we picked individual mid to late L4 hermaphrodites onto seeded plates. One day later, each adult had their average speed measured on an unseeded plate for approximately one minute.

We found that all single mutants were significantly slower than the wild type (Figure 6, *dbl-1(nk3), sma-3(wk30), blmp-1(tm548)*, p<0.0001). There was no significant difference in movement speed between *blmp-1(tm548)* and the double mutants with *dbl-1(nk3)* or *sma-3(wk30)*. These results indicate that the strong *blmp-1(–)* movement defect is epistatic to the moderate movement defect of *dbl-1(–)* and *sma-3(–)* mutants.

**Figure 6.**
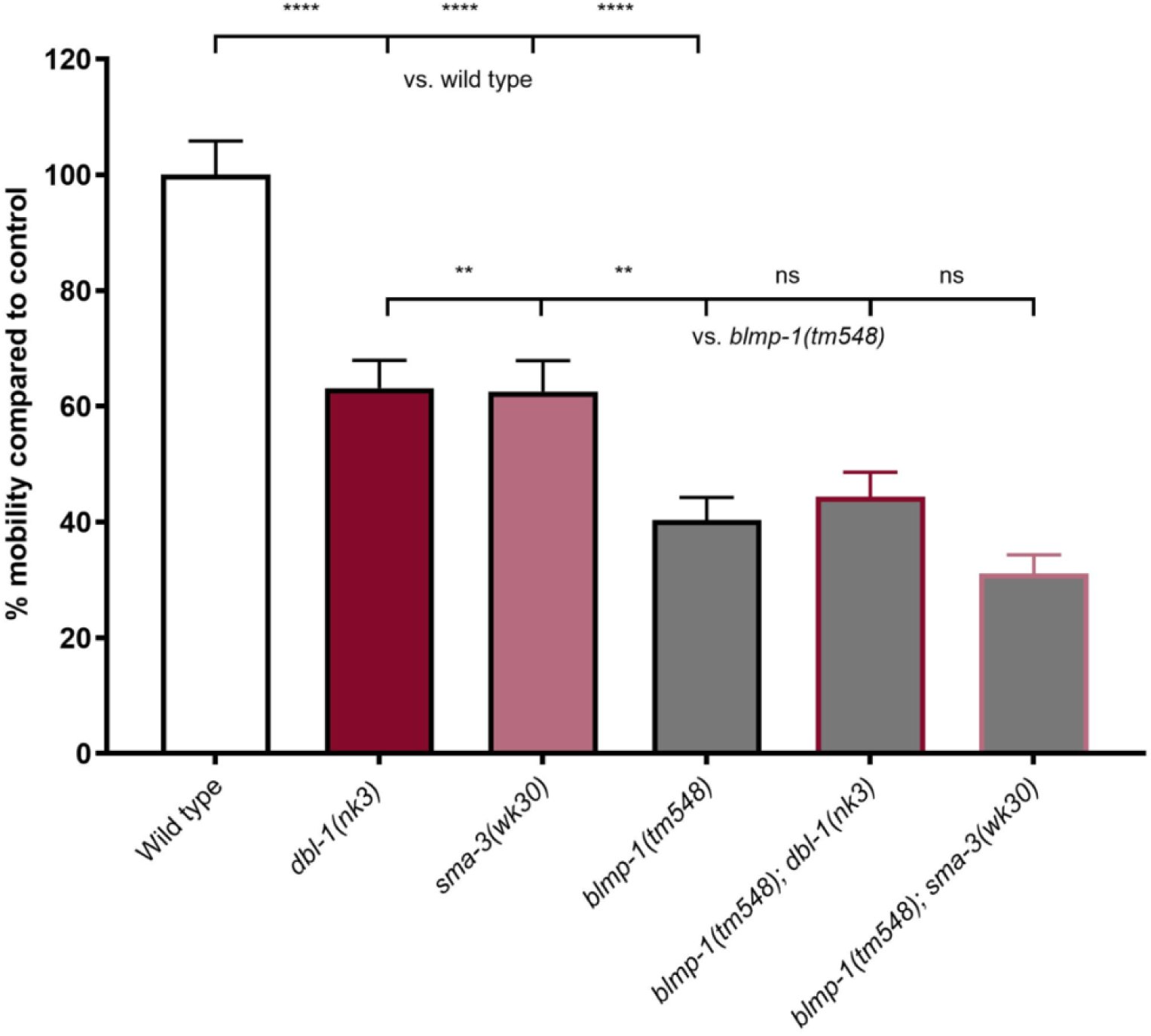
Loss of *dbl-1*, *sma-3*, or *blmp-1* significantly reduces average adult movement compared with wild type animals, with the *blmp-1(tm548)* animals showing the largest reduction. Average speed *per* animal was recorded for each animal one day after the L4-to-adult molt. Data shown is the average of five trials, each with three to twenty animals per strain, normalized to wild type. Error bars represent +/- 1 SEM. ** = <0.01, **** = <0.0001 compared to wild type.

### The *blmp-1(tm548)* allele negatively affects survival post-adult molt

During the movement analysis, we noticed that some animals containing *blmp-1(tm548)* died young, consistent with previously published observations of animals with reduced *blmp-1* expression (Horn et al. 2014; Samuelson et al. 2007a). DBL-1 pathway signaling affects survival of *C. elegans* fed pathogenic bacteria but not *E. coli* OP50 (Mallo et al. 2002; Zugasti and Ewbank, 2009; Tenor and Aballay, 2008; Mørch et al., 2021; Madhu et al. 2023). To quantify survival we followed individual L4s on *E. coli* OP50 from each of the strains used in the movement analysis and scored them for response to harsh touch at 24h, 48h and 72h post mid-L4. Survival was analyzed by Chi-square test (Chi square = 77.8, d.f. = 8, p<0.001) followed by pairwise analysis using the Log-Rank test. We found that defects in *dbl-1* pathway signaling had no effect on survival (Figure 7, *dbl-1* p=0.3, *sma-3* p=1). We found that the *blmp-1(tm548)* strain had reduced survival (p=0.00011 vs. wild type). Neither *dbl-1(nk3)* (p=0.27) nor *sma-3(wk30)* (p=0.83) significantly affected survival of strains with *blmp-1(tm548)* compared to survival of the *blmp-1(tm548)* strain alone. These results indicate that the lifespan reduction caused by loss of BLMP-1/PRDM1 is not further reduced by loss of DBL-1/BMP signaling.

**Figure 7.**
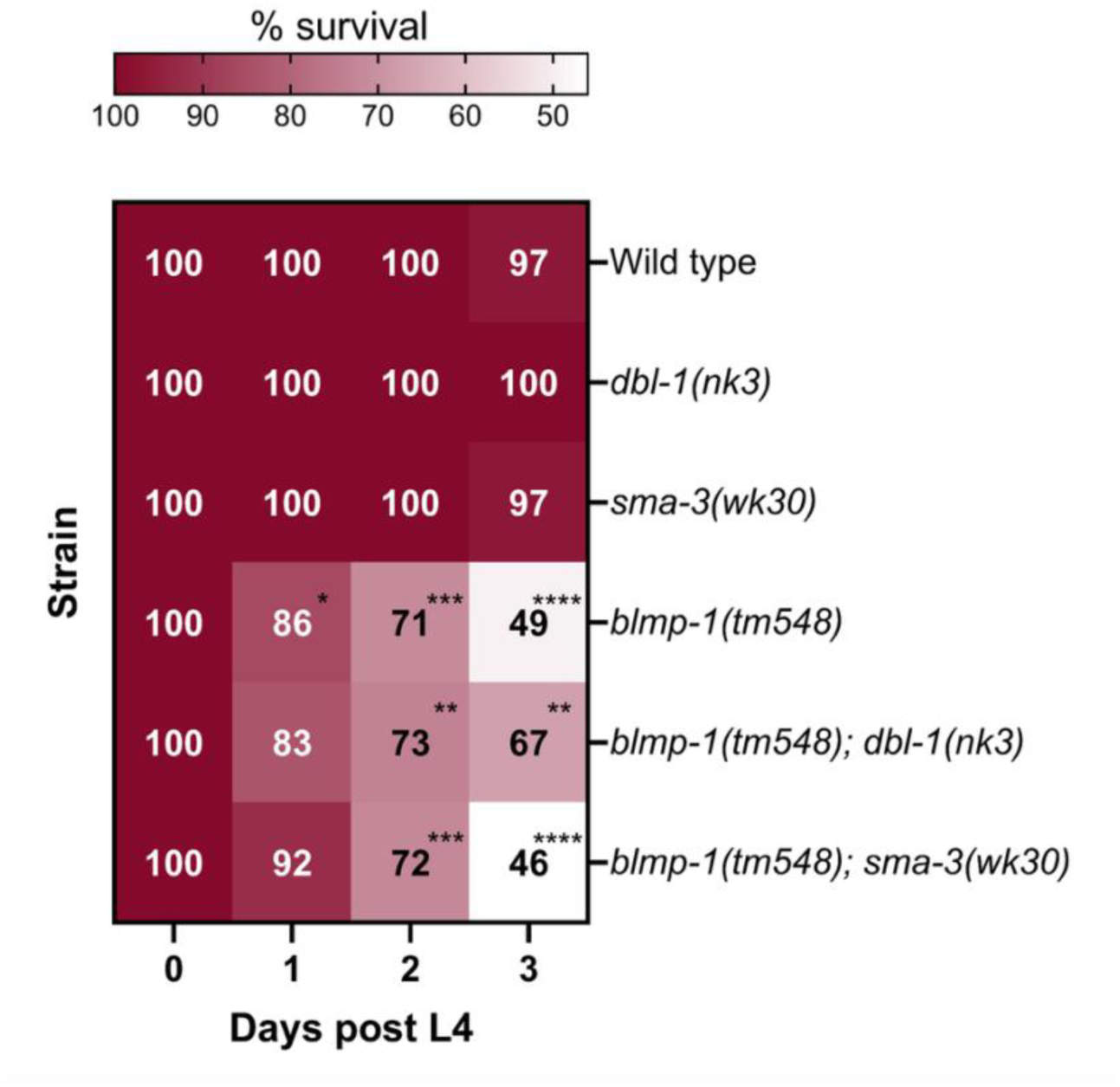
The *blmp-1(tm548)* allele negatively affects survival. Individual L4 animals were transferred to NGM-OP50 plates and scored as alive in response to touch 24, 48 and 72 h later. Between 30 and 50 animals total *per* strain were assayed over three separate trials. * = <0.05, ** = <0.01, **\*\*\*** = <0.001, **** = 0.0001 compared to wild type.

### *blmp-1(–)* suppresses DBL-1/BMP signaling activity

Because our work revealed genetic interactions between the DBL-1/BMP pathway and *blmp-1*, we asked if *blmp-1(–)* impacts DBL-1 pathway target gene regulation. We previously showed that animals lacking BLMP-1/PRDM1 activity upregulate a negative reporter of DBL-1/BMP signaling in adult animals (Lakdawala et al. 2019). However, we later showed that this reporter is responsive to other signaling pathways independent of DBL-1/BMP signaling (Madhu et al. 2020). To validate if *blmp-1(–)* specifically suppresses DBL-1/BMP signaling activity, we generated a strain with *blmp-1(tm548)* and a RAD-SMAD reporter. This reporter is composed of five Smad-binding elements upstream of a GFP coding sequence and is directly regulated by DBL-1/BMP signaling activity, with highest activity at the L2 stage (Tian et al. 2010). Loss of BLMP-1/PRDM1 significantly reduced the GFP reporter activity at the L2 stage compared to RAD-SMAD in a wild-type background, clearly demonstrating that loss of BLMP-1/PRDM1 suppresses DBL-1/BMP signaling activity in L2 (Figure 8).

**Figure 8.**
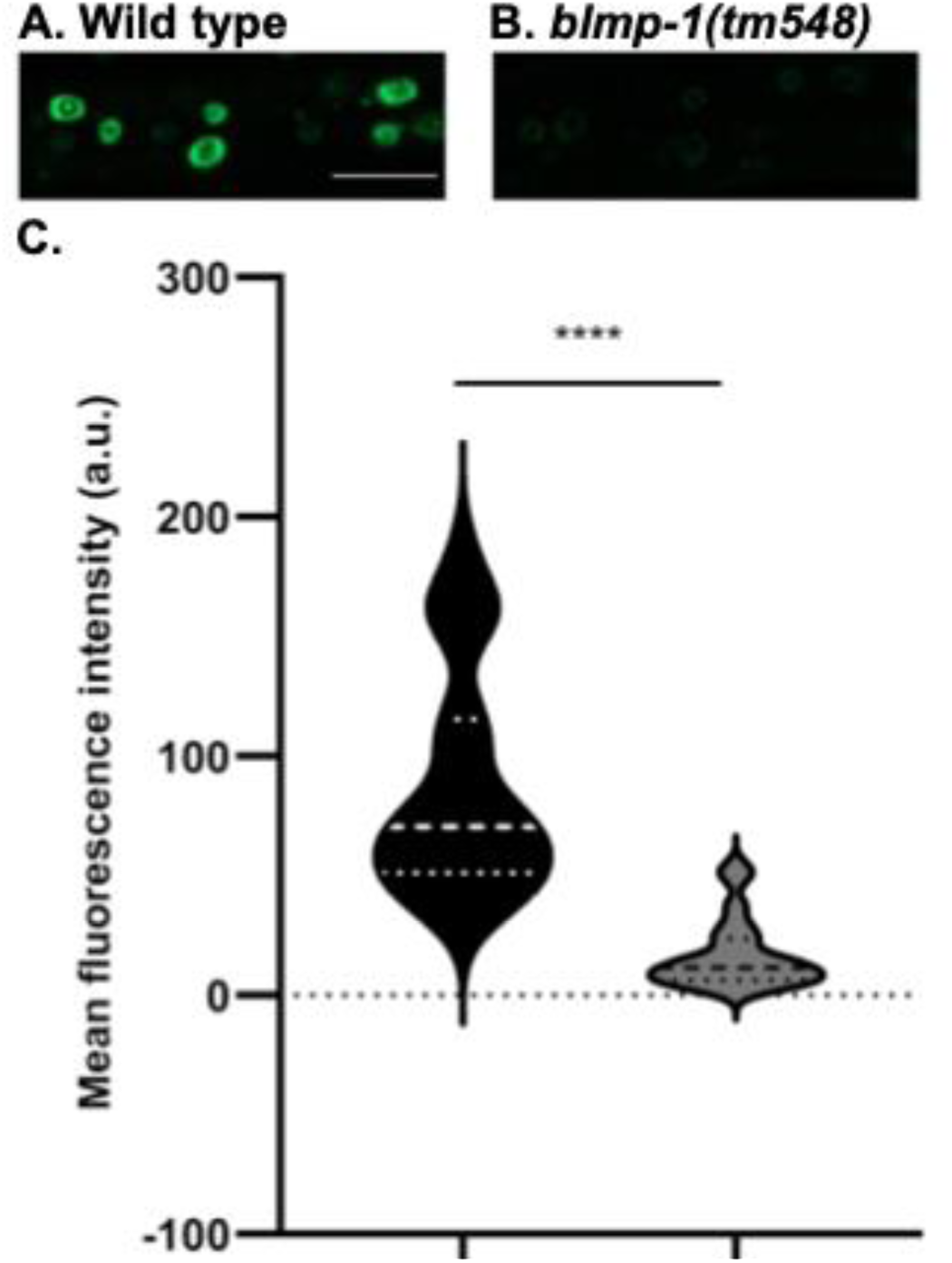
BLMP-1/PRDM1 loss decreases expression of a Smad activity reporter. Animals expressing RAD-SMAD GFP in hypodermal nuclei in a *blmp-1(tm548)* background show decreased fluorescence intensity compared to control. Mean GFP fluorescence intensity of L2 animals expressing the RAD-SMAD reporter in: A, wild-type (control) or B, *blmp-1(tm548)* background was measured at L2 stage. Imaging conditions were consistent between the two groups. Scale bar, 15 µm. C, Mean RAD-SMAD fluorescence intensity of five hypodermal nuclei *per* animal was determined and compared. n = at least 15 animals *per* condition. Error bars represent standard deviation. **** *p* < 0.0001, compared by unpaired *t*-test.

### The DBL-1/BMP signaling pathway regulates *blmp-1* levels at L4 stage

Previous work showed *blmp-1* to be differentially upregulated in adult animals lacking *dbl-1* by RNA sequencing (Lakdawala et al. 2019). To characterize how DBL-1/BMP signaling regulates *blmp-1* expression, we mined existing chromatin ChIP-seq datasets and used bioinformatics tools to identify Smad binding sites upstream of *blmp-1* start site. Published chromatin ChIP-seq datasets of SMA-3 include SMA-3 binding peaks within 1 kb of *blmp-1* (Figure 9A; (Madaan et al. 2018)). Smad proteins bind to specific regions of DNA called Smad Binding Elements (SBE), which have the sequence GTCT or AGAC (Jonk et al. 1998; Madaan et al. 2018). We identified five SBE sequences in the DNA sequence 1 kb upstream of the *blmp-1* start site and three downstream of the start site, consistent with ChIP-seq results (Figure 9B).

**Figure 9.**
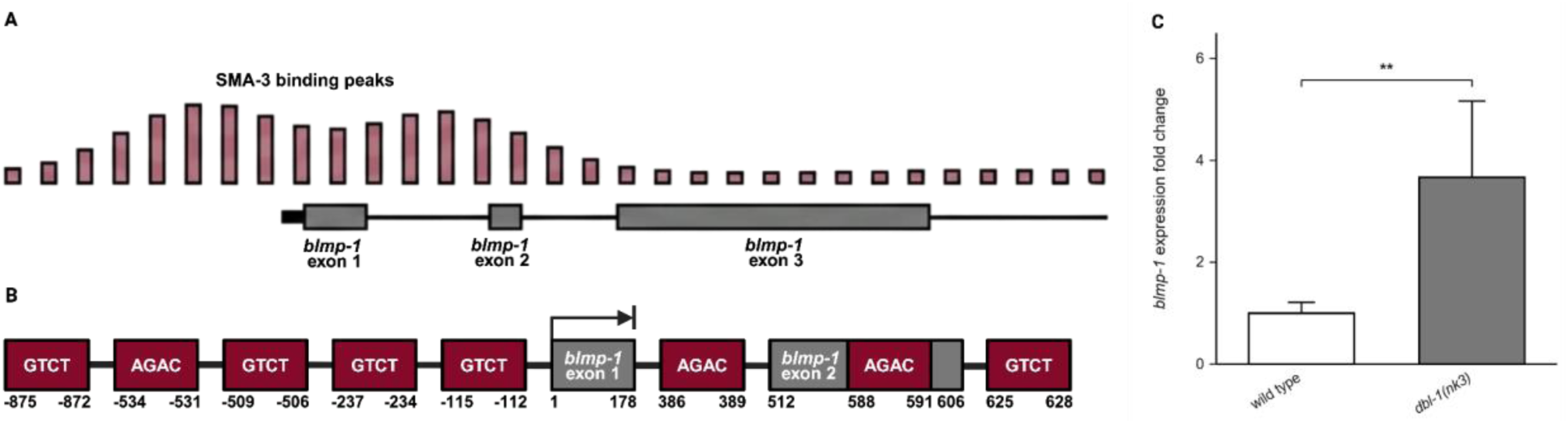
DBL-1/BMP signaling affects *blmp-1* gene expression. **(A)** Adapted from encodeproject.org showing *sma-3* peaks upstream of and within *blmp-1*. (**B)** Graphical representation of SBE upstream of and within the *blmp-1* cDNA. **(C)** Relative mRNA expression levels of *blmp-1* in wild-type control or *dbl-1* mutant animals at the L4 stage were quantified by real-time PCR. The experiment was performed in four biological replicates with three technical replicates each. Error bars represent standard error of the mean. ** *p* < 0.01, compared by unpaired *t*-test.

To determine if DBL-1/BMP signaling regulates *blmp-1* expression in L4, when BLMP-1/PRDM1 is required for development of adult structures, we measured *blmp-1* expression levels using qRT-PCR at the L4 stage in *dbl-1* mutants and compared them to WT control levels. A significant increase, averaging a 3.7-fold difference, of *blmp-1* levels in *dbl-1* mutant animals compared to WT was observed (Figure 9C). Collectively, the ChIP-seq and qRT-PCR analyses indicate DBL-1/BMP signaling pathway directly suppresses *blmp-1* gene expression via SMA-3.

### BLMP-1/PRDM1 physically interacts with Smads to regulate common target genes

We next asked if *blmp-1* is not just an important target gene of the DBL-1/BMP pathway but is also a Smad binding partner. Smads and BLMP-1/PRDM1 both interact with other proteins in a larger protein complex to selectively bind to DNA and regulate gene expression (Minnich et al. 2016; Conidi et al. 2011). These protein interactions are dynamic and context dependent. However, Smads and BLMP-1/PRDM1 have not been shown to interact in the same complex. To determine if BLMP-1/PRDM1 and Smads interact, we performed yeast-two hybrid analyses with BLMP-1/PRDM1 as bait and SMA-2, SMA-3, and SMA-4 as prey (Fields and Song 1989). No growth was seen in the auto-activation controls. Growth was seen for yeast co-transformed with BLMP-1-BD and each of the three SMA-AD plasmids, indicating a physical interaction between BLMP-1/PRDM1 and all three SMA proteins (Figure 10).

**Figure 10.**
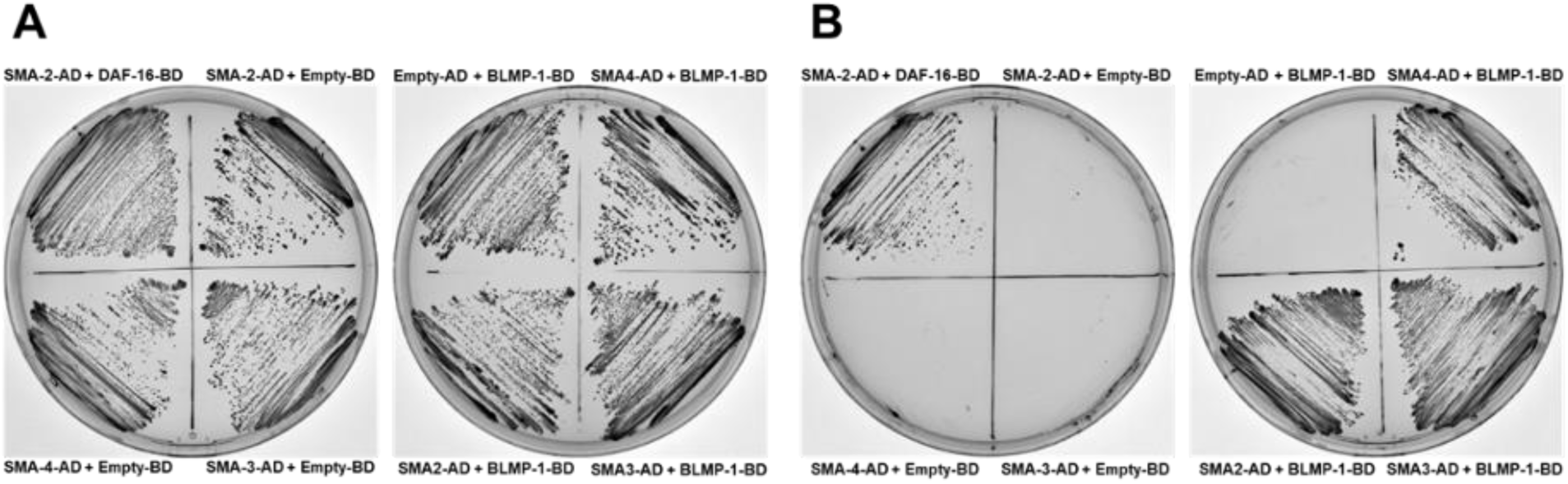
BLMP-1/PRDM1 physically interacts with Smads. Yeast were co-transformed with BLMP-1-binding domain (BD), SMA-activation domain (AD) proteins, and controls. **(A)** Control plates with –Leu, –Trp yeast media to grow all co-transformants. Successful co-transformation of SMA-2-AD, SMA-3-AD, and SMA-4-AD with empty or DAF-16-BD plasmid (left) and BLMP-1-BD with SMA-2-AD, SMA-3-AD, and SMA-4-AD, or empty AD plasmid, were co-transformed as controls for auto-activation. DAF-16-BD co-transformed with SMA-2-AD was used as a positive control (Qi et al., 2017). **(B)** Screen plates with selective –Ade, –His, –Leu, and –Trp yeast media. Growth was seen in the positive control with SMA-2-AD and DAF-16-BD.

We hypothesized that BLMP-1/PRDM1 and SMA-3/R-Smad control the expression of an overlapping set of target genes that control shared traits because they have genetic interactions and physically interact with each other. To interrogate this hypothesis, publicly available ChIP-seq datasets of SMA-3/R-Smad and BLMP-1/PRDM1 were compared (Table S3, (Gerstein et al. 2010)). SMA-3/R-Smad has 4,204 binding regions in the genome whereas BLMP-1/PRDM1 has 3,030 binding regions. We identified the binding regions of SMA-3/R-Smad and BLMP-1/PRDM1 that have at least one base pair overlap. We found 1,958 regions where SMA-3/R-Smad and BLMP-1/PRDM1 binding peaks intersected with each other (47% of SMA-3/R-Smad binding regions and 65% of BLMP-1/PRDM1 binding regions are shared). Approximately 95% of these overlapping regions are in gene promoter regions (Figure 11, Table S3). These results indicate that SMA-3/R-Smad and BLMP-1/PRDM1 not only interact with each other but also regulate common target genes.

**Figure 11.**
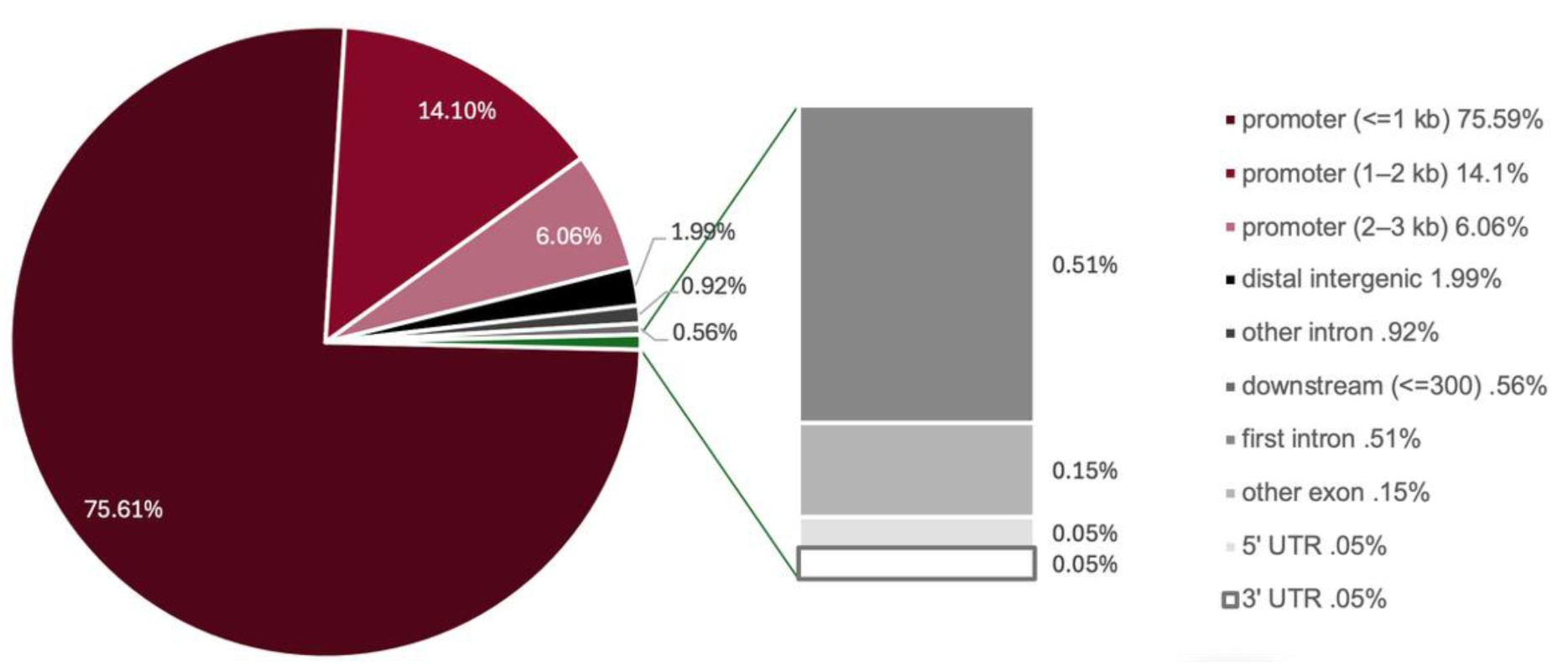
SMA-3/R-Smad and BLMP-1/PRDM1 bind common chromosomal sites primarily within 2 kb of target gene start sites. Genomic annotation of ChIP-seq peaks common between SMA-3/R-Smad and BLMP-1/PRDM1.

To further shortlist genes that are regulated by DBL-1/BMP pathway and BLMP-1/PRDM1 together, previously published differential gene expression datasets of genes that are differentially regulated by BLMP-1/PRDM1 or DBL-1/BMP pathway components were compared (Roberts et al. 2010; Madhu et al. 2023; Luo et al. 2010; Liang et al. 2007; Huang et al. 2014; Horn et al. 2014). Many common genes that are differentially regulated by BLMP-1/PRDM1 or DBL-1/BMP signaling were identified. To further validate this result, we sampled signaling and structural genes that may have roles in traits determined by BLMP-1/PRDM1 and DBL-1/BMP pathway and are either upregulated or downregulated in both DBL-1/BMP pathway mutants and *blmp-1* mutant animals. We compared expression levels of these shortlisted genes in wild-type, *blmp-1(tm548)*, *dbl-1(–)*, and double mutants by qRT-PCR and found *grd-6* and *col-117* levels are significantly downregulated in *dbl-1* and *blmp-1* single mutants as well as in *blmp-1(tm548); dbl-1(–)* double mutants (Figure 12A, B). Conversely, *wrt-1*, and *col-128* levels were found to be significantly upregulated in single mutants as well as double mutants (Figure 12C, D). These results indicate DBL-1/BMP signaling and BLMP-1/PRDM1 regulate expression of common genes. Overall, these results from bioinformatics analyses, yeast two-hybrid, and qRT-PCR assays indicate that BLMP-1/PRDM1 physically interacts with the Smads to regulate expression of common target genes.

**Figure 12.**
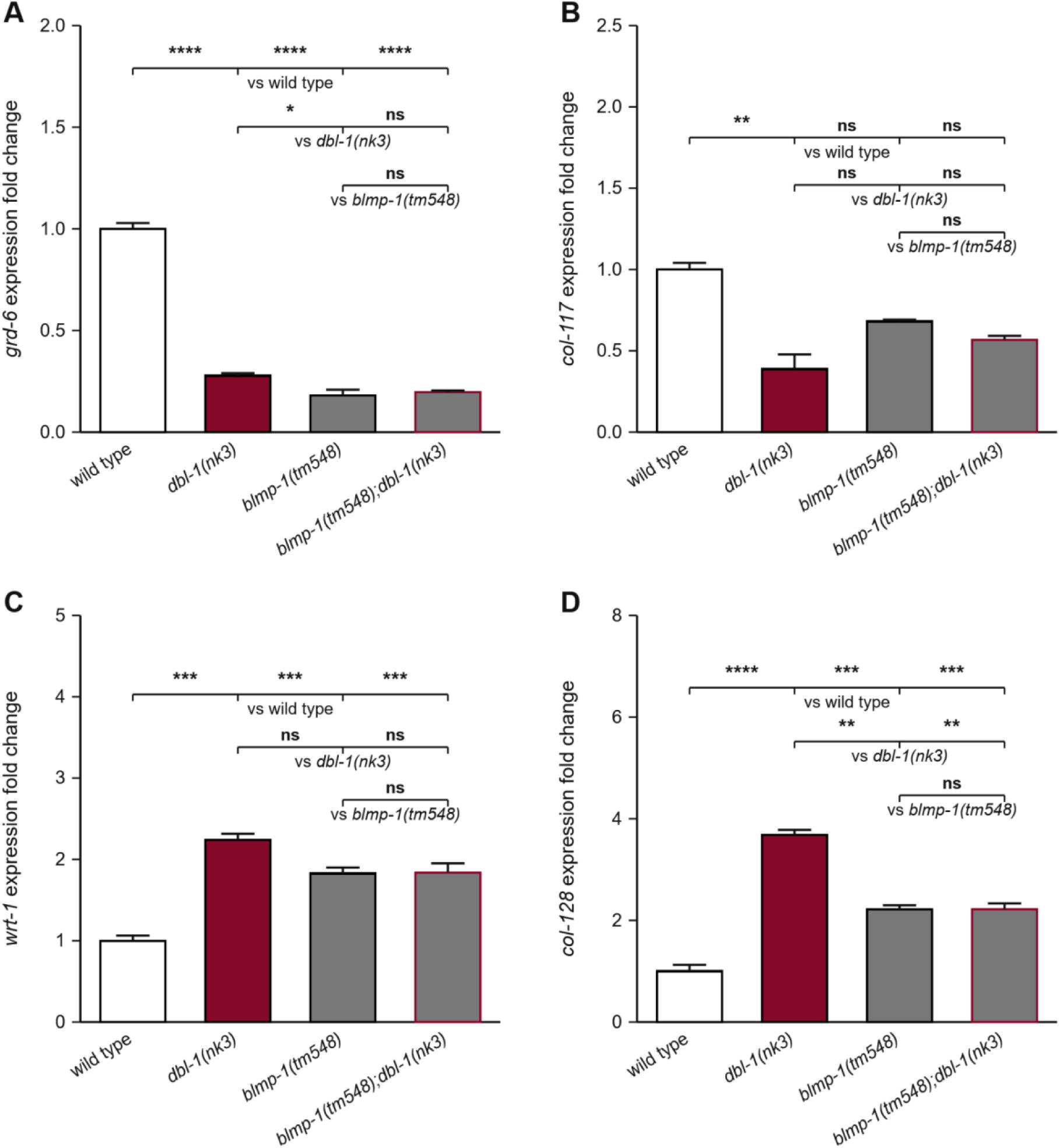
The DBL-1/BMP signaling pathway and BLMP-1/PRDM1 regulate common genes. Relative mRNA expression levels of **(A)** *grd-6*, **(B)** *col-117*, **(C)** *wrt-1* and **(D)** *col-128* were quantified by real-time PCR in wild-type control, *dbl-1(–)*, *blmp-1(tm548)*, and *dbl-1(–); blmp-1(tm548)* animals at L2 stage. **(A, B)** mRNA expression of *grd-6* and *col-117* is reduced in *dbl-1*, *blmp-1*, and *blmp-1; dbl-1* mutant background as compared to control. **(C, D)** mRNA expression of *wrt-1* and *col-128* is increased in *dbl-1*, *blmp-1* and *blmp-1; dbl-1* mutant background as compared to control. Experiments were performed in three technical replicates. Error bars represent standard error mean. ** *p* < 0.005, **** *p* < 0.0005, **** *p* < 0.0001, compared by one-way ANOVA using Tukey’s multiple comparisons test.

## Discussion

Here, we identify how a conserved *C. elegans* TGF-β superfamily signaling pathway may spatiotemporally coordinate expression of different genes that are important for proper development of multiple traits by its interactions with pioneer transcription regulator BLMP-1/PRDM. We previously showed that DBL-1/BMP signaling genes genetically interact with BLMP-1/PRDM1 in *C. elegans* (Lakdawala et al. 2019). We have expanded their genetic interactions and identified novel cellular and molecular interactions between DBL-1/BMP signaling and BLMP-1/PRDM1 (Figure 13). In addition to genetic and cellular interactions, we identified a physical interaction between the DBL-1/BMP pathway Smads and BLMP-1/PRDM1. This physical interaction is likely meaningful because, at L2 at least, almost 2000 target genes of SMA-3/R-Smad and BLMP-1/PRDM1 are shared, which is about half of all SMA-3/R-Smad target genes and two thirds of BLMP-1/PRDM1 target genes. Our analyses suggest that BLMP-1/PRDM1, in its gatekeeper role, first helps remodel genomic regions containing Smad target sequences and then recruits Smads to regulate expression of thousands of target genes and resulting traits throughout larval development and reproduction. We propose that this Smad-PRDM axis is an ancestral mechanism for metazoan development and diversification.

**Figure 13.**
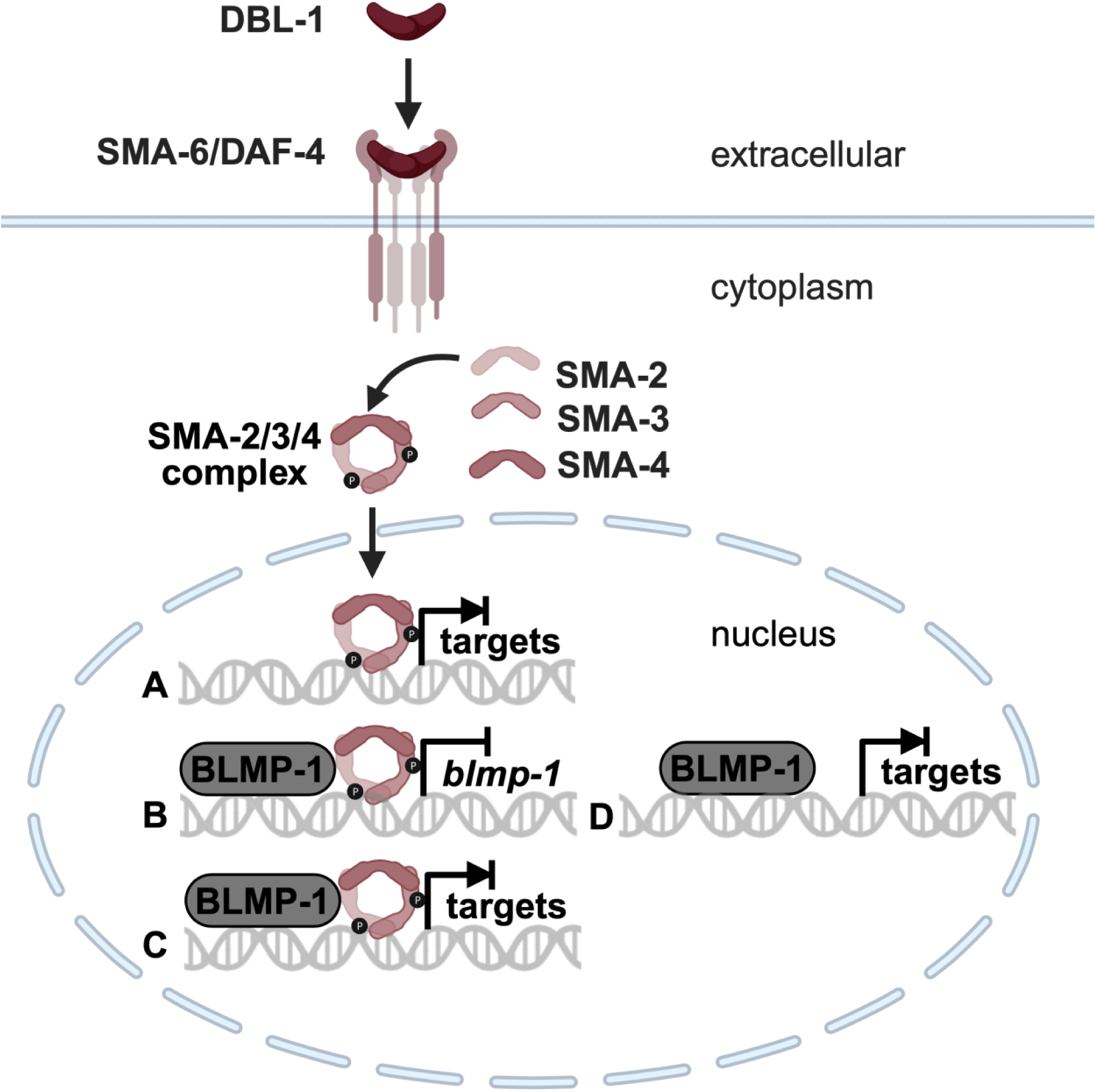
Model of SMAD complex-BLMP-1/PRDM1 interactions driving expression of different sets of target genes. The SMAD complex regulates gene transcription independent of BLMP-1, possibly with a different (not shown) pioneer transcription factor (A). BLMP-1, by homology and consistent with published genetic analyses and this work, is proposed to act as a pioneer factor. This work showed that BLMP-1 binds SMA-2, SMA-3, and SMA-4. The SMAD complex and BLMP-1 co-regulate thousands of target genes (B and C), including *blmp-1* (B). BLMP-1 can also work without SMADs, possibly with other transcription factors (not shown), for SMAD-independent gene regulation (D).

### Coordination of multiple traits by DBL-1/BMP pathway signaling and BLMP-1/PRDM1

DBL-1/BMP and BLMP-1/PRDM1 regulate many shared processes (Table S1) (Zhang et al. 2012; Savage-Dunn and Padgett 2017; Nelson et al. 2011; Gumienny and Savage-Dunn 2013). In humans, too, there are many striking phenotypic parallels between TGF-β superfamily signaling and PRDM1 in tissue patterning and development, including in digit formation, craniofacial development, and skin differentiation (Vincent et al. 2005; Robertson et al. 2007). The interaction of BLMP-1/PRDM1 with the DBL-1/BMP pathway was first shown when loss of *blmp-1* suppressed the long body size of animals with increased DBL-1/BMP signaling (Lakdawala et al. 2019). Here, we provide support that this genetic interaction is additive for body size because animals lacking BLMP-1/PRDM1 and DBL-1/BMP signaling are smaller than animals with single mutations in either *blmp-1* or the DBL-1/BMP pathway (Figure 1). This result suggests that either *blmp-1* and DBL-1/BMP signaling work independently to regulate genes involved in body size determination or that they regulate shared target genes as well as body size genes that are responsive to only *blmp-1* or DBL-1/BMP signaling. We favor the latter hypothesis because body size is a multifactorial trait and many genes identified in these three categories (SMA-3 only, BLMP-1 only, and both) may impact body size (Table S3).

However, for most phenotypes analyzed, *blmp-1(tm548)* masked the DBL-1/BMP pathway mutant traits. For male tail morphogenesis, the double mutants had complete penetrance of the *blmp-1(lf)*-associated over-retraction (Ore) defect (Figure 2). This finding is consistent with the precocious male tail morphogenesis caused by loss of BLMP-1/PRDM1 preventing the delayed migration (Lep) phenotype seen in animals lacking DBL-1/BMP signaling from occurring (Nelson et al. 2011). BLMP-1/PRDM1 is needed for proper migration of the developing hermaphrodite gonad during L3 and L4 stages. Loss of DBL-1/BMP exacerbates the penetrance of the migration defect in populations of animals with a hypomorphic allele of the *unc-5* netrin receptor (Merz et al. 2003). However, loss of DBL-1/BMP signaling pathway components did not further increase the percentage of *blmp-1(lf)* animals with mis-migrated gonads (Table 1). *blmp-1* also showed an epistatic relationship with DBL-1/BMP pathway mutants in L4s/young adults for lipid content and in one-day old adults with the movement defects (Figures 5, 6). Together, these genetic analyses support a model in which BLMP-1/PRDM1 is required (in whole or in part) for DBL-1/BMP pathway Smads to function throughout the lifespan of the organism.

A previous report noted a reduction of lipid levels in the anterior intestine in both *dbl-1* and *sma-3* mutants (Clark et al. 2021). We found that the lack of SMA-3/R-Smad and BLMP-1/PRDM1 reduced whole animal lipid levels, but were not additive, suggesting that the two work together to influence lipid levels (Figure 5). Lack of DBL-1/BMP had no effect on whole animal lipid levels, most likely due to methodological differences. As the intestine is the main site of lipid synthesis and storage in *C. elegans*, measuring lipid levels at the whole animal level is most likely less sensitive than restricting the analysis to the intestine alone, a hypothesis supported by our reporting of a smaller decrease in lipid levels in *sma-3* animals than previously measured.

Our findings for brood size suggest there may be a difference in how DBL-1/BMP and SMA-3/R-Smad affect this trait in animals lacking BLMP-1/PRDM1 (Figure 4). Defects in the germline or somatic gonad can lead to reduced brood size. DBL-1/BMP signaling pathway plays an important role in germline development, including in oocyte quality maintenance and germline proliferation (Luo et al. 2009; Luo et al. 2010; Huang et al. 2014; Horn et al. 2014). The larval function of BLMP-1/PRDM1 is critical for proper gonad migration, which may secondarily affect brood size. In this study, *blmp-1* was epistatic to *dbl-1* in progeny number, but the *blmp-1; sma-3* strain had a lower brood size than either single mutant. This observation suggests that the Smads may crosstalk with other pathways that also impact brood size. In other organisms, crosstalk occurs between Smads and Notch, Wnt, and other conserved signaling pathways, so these interactions may also exist in *C. elegans* (Luo 2017; Derynck et al. 2014).

### DBL-1/BMP signaling transcriptionally regulates *blmp-1* expression

In *C. elegans*, the BLMP-1/PRDM1 expression pattern is highly dynamic during development, however not much is known about its upstream regulation (Stec et al. 2021; Huang et al. 2014; Horn et al. 2014). Bioinformatics analyses reveal potential SMA-3/R-Smad binding sites upstream of *blmp-1* and ChIP-seq data showed SMA-3 binding at the *blmp-1* locus, suggesting *blmp-1* levels are directly regulated during postembryonic development by the DBL-1/BMP pathway (Figure 9A, B, (Lakdawala et al. 2019; Gerstein et al. 2010). We previously showed that *blmp-1* is transcriptionally regulated by DBL-1/BMP signaling during postembryonic development (Lakdawala et al. 2019). This work expands on that interaction, showing that DBL-1/BMP signaling represses *blmp-1* expression (Figure 9C). Previous work in other vertebrate and invertebrate systems showed that BMP positively regulates PRDM1/BLIMP1, PRDM14, and PRDM16 expression during embryonic development (Yamaji et al. 2008; Saitou et al. 2005; Ohinata et al. 2005; Hopf et al. 2011; He et al. 2025). *Prdm16* has Smad binding sites that are bound by pSMAD1/5/8 but not pSMAD3 in response to BMP4 (He et al. 2025). However, it is unclear if BMP4’s regulation of PRDM1 or PRDM14 is direct. This work provides evidence that the DBL-1/BMP pathway plays a key role in controlling the dynamic expression pattern of BLMP-1/PRDM1 during postembryonic development in *C. elegans*.

### DBL-1/BMP pathway Smads physically interact with BLMP-1/PRDM1

Smad complexes bind DNA rather poorly and must interact with other transcriptional regulators to bind DNA and regulate transcription of target genes (Morikawa et al. 2011; Hill 2016). Smads bind to chromatin remodeling proteins and to the Mediator complex that brings in RNA polymerase I to transcribe target genes. In *C. elegans*, the Smad machinery interacts with insulin-like signaling pathway component DAF-16/FOXO to regulate germline proliferation (Qi et al. 2017). Similarly, BLMP-1/PRDM1 can have different binding partners to regulate different processes (Hyun et al. 2016; Fong et al. 2020). PRDM3/16 subfamily members are the only PRDM proteins in mammals known to interact with BMP ligands (Warner et al. 2007; Takahata et al. 2009; Sato et al. 2008; Hohenauer and Moore 2012; Bjork et al. 2010; Alliston et al. 2005). Our work revealed a novel physical interaction of the Smad machinery with BLMP-1/PRDM1. Moreover, analyses of previously published datasets as well as qPCR analyses show Smads and BLMP-1/PRDM1 have an extensively overlapping set of target genes. PRDM1/Blimp1 in mice and BLMP-1/PRDM1 in *C. elegans* are known to interact with chromatin remodeling components and to prime DNA for additional transcription factors to bind, thereby increasing their transcriptional output (Stec et al. 2021; Nadeau and Martins 2021; Fong et al. 2020). Our work suggests that BLMP-1/PRDM1 proteins remodel Smad target sequences and then recruit Smads for transcription as part of a larger transcription complex. Without BLMP-1/PRDM1, the Smads may not effectively bind to the shared target gene sites and control their expression. In this way, loss of BLMP-1/PRDM1 masks the loss of a Smad or DBL-1/BMP signaling.

Smad interactions with different proteins lead to diverse functional outputs (Conidi et al. 2011). Differences in the traits caused by loss of DBL-1/BMP pathway function and loss of BLMP-1 function may occur because the Smad complex and BLMP-1 can also interact with other proteins to regulate transcription of shared target genes differently. Differences may also occur because sets of target genes that are responsive to either Smads or BLMP-1/PRDM1, but not both, may account for the phenotype differences.

### Evolutionary implications of a conserved Smad-PRDM transcriptional module

BMP pathway components and PRDM family members are found in metazoans from sponges to mammals. These gene families experienced diversification and losses as speciations occurred, which may have helped drive metazoan diversification (Vervoort et al. 2016). Combined with phylogenetic studies, our work in C. *elegans* suggests that the interaction between Smads and PRDM superfamily proteins first identified in vertebrate systems is an ancient mechanism for regulating BMP signaling and myriad traits that distinguish species. While the physical interaction between vertebrate Smads and PRDM3/16 members is well documented, our BLMP-1/PRDM1 findings hint that the genetic interactions seen between vertebrate TGF-β pathway genes and PRDM1 genes may also be caused by physical associations between Smads and PRDM1 members. It will be of interest to see if this is the case.

Our work studied the interaction of Smads and BLMP-1/PRDM1 for traits associated with non-stressed development, reproduction, and aging. It will also be of interest to determine how DBL-1/BMP and BLMP-1/PRDM1 interact to regulate responses in changing stress and non-stress environments. DBL-1/BMP signaling is important for normal larval development and reproduction. Loss of *dbl-1* function increases the incidence of dauer formation of animals lacking the dauer-inhibiting DAF-7 TGF-β superfamily member at 20°C, showing that DBL-1 normally helps prevent animals from entering the dauer pathway (Morita et al. 1999). In addition, the *dbl-1* gene is down-regulated in dauers, indicating DBL-1 activity is not needed as much in dauers (Liu et al. 2004). DBL-1 prevents nuclear localization (and therefore activity) of DAF-16/FOXO, a dauer-inducing transcription factor (Clark et al. 2021). BLMP-1, on the other hand, is required not only for traits formed during reproductive development and adulthood, including traits assayed in this study, it is also needed for proper entry into the dauer diapause state (Hyun et al. 2016; Horn et al. 2014). The conserved signaling circuits that regulate timing of developmental events in *C. elegans*, including the dauer entry stress response, have functions in mammalian stress responses and stem cell fate determination. Exploring the Smad-PRDM axis in this developmental circuit may help illuminate how these complex processes adapt to environmental changes in this roundworm and in other animals (Batlle and Massague 2019; Bikoff et al. 2009; Chandiran et al. 2022; Hohenauer and Moore 2012; Leszczynski et al. 2020; Setiawan et al. 2022).

### Conclusions

Our work provides evidence for a new molecular mechanism by which TGF-β signaling regulates organismal traits. We identified and characterized novel interactions between DBL-1/BMP signaling and a pioneer transcriptional regulator, BLMP-1/PRDM1, in the context of post-embryonic development in *C. elegans*. Our study showed that genetic interactions between vertebrate TGF-β signaling pathways and BLMP-1/PRDM1 homologs are conserved for the DBL-1/BMP pathway in *C. elegans*. We also identified BLMP-1/PRDM1 as a new binding partner of the Smad machinery. BLMP-1/PRDM1 is important for regulating downstream gene targets of the DBL-1/BMP pathway. Collectively, our work adds a new layer of complexity in the regulation of the DBL-1/BMP signaling pathway, revealing that the master chromatin regulator BLMP-1/PRDM1 acts as a pivotal gatekeeper for DBL-1/BMP signaling in *C. elegans*.

Together, our findings support a model that helps explain how multiple traits are determined at different times by DBL-1/BMP. *blmp-1* and the DBL-1/BMP signaling pathway interact with each other at two levels. First, BLMP-1/PRDM1, as the gatekeeper pioneer transcription factor, recruits Smad complexes to target gene sites to inhibit or promote expression of shared target genes and specific traits. Second, DBL-1 signaling can inhibit *blmp-1* expression levels. The transcriptional suppression of *blmp-1* by DBL-1/BMP and the traits affected by both DBL-1 and BLMP-1/PRDM1 suggest an elegant feedback loop that maintains signaling homeostasis in reproductively favorable environments. Therefore, we propose that these two players are parts of a developmental rheostat that is precisely regulated at different times in different tissues to craft the developing animal and control reproduction and survival. Genetic on/off switches are critical for normal development and variations in when they are turned up or down can cause disorders — or new species. Given the deep conservation of TGF-β superfamily signaling components and PRDM family members across the metazoan lineage, discovering if the Smad-PRDM axis in *C. elegans* is conserved in other animals will be of keen interest for understanding these fundamental processes in development, disorders, and evolution.

## Data availability

Strains and other materials are available upon request. The authors affirm that all data necessary for confirming the conclusions of the article are present within the article, figures, and tables. Supplemental material available at GENETICS online.

## Acknowledgements

Christopher Hammell provided *blmp-1* and control constructs for yeast two-hybrid analyses. Dr. Omar Darwish and Dr. Micah Thornton provided bioinformatics and figure assistance. Dr. Thomas Guffey and Dr. Wanyi Wang provided statistical analysis and figure assistance. Some *C. elegans* strains were obtained from the *Caenorhabditis* Genetics Center (CGC), which is funded by NIH Office of Research Infrastructure Programs (P40 OD010440). WormBase (version: WS295), The Alliance of Genome Resources, and The Gene Ontology Knowledgebase were valuable resources. Some figures were created using BioRender. This work is dedicated to family and friends for their constant motivation and support.

